# *ST6GAL1*-mediated heterogeneity in sialic acid levels potentiates invasion of breast cancer epithelia

**DOI:** 10.1101/2020.04.28.065573

**Authors:** Dharma Pally, Durjay Pramanik, Shahid Hussain, Shreya Verma, Anagha Srinivas, Rekha V Kumar, Ramray Bhat

**Affiliations:** Department of Molecular reproduction, Development and Genetics, Indian Institute of Science, Bangalore, India-560012.; Department of Pathology, Kidwai Memorial Institute of Oncology, Bangalore, India-560029.

**Keywords:** Tumor heterogeneity, Sialic acid, Breast cancer, Extracellular matrix, Cancer invasion, Cancer model

## Abstract

Heterogeneity in phenotypes of malignantly transformed cells and aberrant glycan expression on their surface are two prominent hallmarks of cancers that have hitherto not been linked to each other. In this paper, we identify heterogeneity in a specific glycan linkage: α2,6-linked sialic acids within breast cancer cells *in vivo* and in culture. Upon sorting out two populations with moderate and relatively higher cell surface α2,6-linked sialic acid levels from the triple negative breast cancer cell line MDA-MB-231, both populations (denoted as medium and high-2,6-Sial cells respectively) stably retained their levels in early passages. Upon continuous culturing, medium 2,6-Sial cells recapitulated the heterogeneity of the unsorted line whereas high 2,6-Sial cells showed no such tendency. Compared with the high 2,6-Sial, the medium 2,6-Sial cells showed greater adhesion to reconstituted extracellular matrices (ECM) as well as invaded faster as single cells. The level of α2,6-linked sialic acids in the two sublines was found to be consistent with the expression of a specific glycosyl transferase, *ST6GAL1*. Stably knocking down *ST6GAL1* in the high 2,6-Sial cells, enhanced their invasiveness. When cultured together, medium 2,6-Sial cells differentially migrated to the edge of growing tumoroid-like cultures, whereas high 2,6-Sial cells formed the central bulk. Simulations in a Cellular Potts model-based computational environment that is calibrated to our experimental findings suggest that the heterogeneity of cell-ECM adhesion, likely regulated by α2,6-linked sialic acids facilitates niches of highly invasive cells to efficiently migrate centrifugally as the invasive front of a malignant tumor.

**Significance Statement:** Cell-surface sugars are aberrantly expressed in cancer but their contributions to tumor heterogeneity are not known. In this study, we uncover and separate breast cancer populations with distinct α2,6-linked sialic acid levels. The moderately expressing population shows stronger adhesion to extracellular matrix than the high expressing population. It also invades faster through the matrix as single cells. Combining experiments with computational modelling, we show that the heterogeneity in matrix adhesion is vital to accentuating cell invasion. In some conditions, invasion of heterogeneous populations may compare with, or exceed that of, homogeneous moderately expressing populations. Our findings are vital to furthering our understanding of how cancers spread and potentially qualify efforts to manage the disease through glycan-editing or immunotherapeutic approaches.

## Introduction

One of the hallmarks of malignant tumors is the heterogeneity in the phenotypes of its constituent transformed epithelia. Observations of phenotypic heterogeneity can be traced back to the demonstration by Hawkins and co-workers, of variable expression of estrogen receptor (ER) among cells within a single tumor. With time, evidence of intratumoral variation in expression was discovered for several genes/markers and is responsible for determining clinical behaviour and response to treatment (1–5). Intratumoral heterogeneity can also contribute to misdiagnosis of the aggressiveness and grade of breast cancer leading to its mistreatment (6–8).

A combination of genomic and epigenomic aberrations, loss in a dynamic and reciprocal regulation of homeostasis by the tissue microenvironment and stochasticity leads to diversity in protein expression, localization and interaction within cells belonging to the same population. This diversity in turn leads to heterogeneity in cellular phenotypes (9–12). However, proteins are not the only molecular species to show such alterations in malignant contexts. Changes in levels of sugars on the surfaces of cancer cells, when compared with their untransformed counterparts have been demonstrated since a long time (13–15). Further studies show that altered levels of N- and O-linked glycosylations in transformed epithelia as well as in tumor associated stromal cells impact the progression and metastasis of cancer as well as its response to chemotherapy (16–18). For example, aberrantly glycosylated β1-integrin leads to altered cell-ECM adhesion thereby aiding cancer cell invasion and metastasis (19). An increase in Sialyl Lewis^X^ leads to increased adhesion of cancer cells to endothelial cells via selectin leading to colonization of distant organs (20). O-GlcNAylated c-Myc can compete with its unglycosylated counterpart in phosphorylation leading to increased stability and thereby increase cancer cell proliferation (21). Hypersialylation is one of the most frequently observed changes in glycosylation in many cancer types (22, 23). Selective enrichment of terminal α2,6-linked sialic acids (referred here-onward as α2,6-Sial), due to overexpression of *ST6GAL1*, in cancer cell glycocalyces can elicit a wide range of biological outcomes like protection from hypoxia, resistance to chemotherapy, pro-survival and conferral of cancer stem cell phenotype (24–27).

To the best of our knowledge, there is no literature on heterogeneous levels of glycans on the surface of transformed cells within growing tumors. In the present study, we investigated this question in the context of sialic acid expression in breast cancer. Using a combination of lectin-based flow cytometry and cytochemistry, we demonstrated diverse expression profiles of α2,6-Sial within breast cancer cells in vitro and *in vivo*. The diversity was glycan-specific: α2,3-Sial or other assessed oligosaccharides did not show any such heterogeneous expression. We found two distinct cell populations with moderate and high levels of α2,6-Sial in the triple negative breast cancer cell line MDA-MB-231. Combining cell biological assays with agent-based modelling simulations, we demonstrate how the reported intercellular glycan heterogeneity results in the differential migration of more invasive epithelia to the invading edge of cultured 3D tumoroids. A better understanding of intratumor glycobiological heterogeneity is certain to impact breast cancer diagnosis and treatment in the future.

## Results

### Breast cancer epithelia show heterogeneity in α2,6-Sial levels

We assayed for intercellular differences in α2,6- and α2,3-Sial levels in breast cancer sections using lectin cytochemistry. (FITC-conjugated *Sambucus nigr*a (SNA) lectin and TRITC-conjugated *Maackia amurensis* (MAA) lectin were used as probes for α2,6- and α2,3-Sial respectively). Tumor sections showed signals for both sugar linkages when compared to negative sugar controls (Fig S1&1). However, although cells in the sections stained uniformly for α2,3-Sial (Fig 1A, red), cellular staining for α2,6-Sial (Fig 1A, green) was variegated: rounded patches of cells with high levels of α2,6-Sial were surrounded by dispersed populations with comparatively lower levels (Fig 1A). This was confirmed through per-cell quantification of cancer cells that revealed a greater variance in cell-specific expression of α2,6-Sial relative to α2,3-Sial (Fig 1B). Whereas elevated levels of α2,6-Sial in breast cancer epithelia have been previously reported (28, 29), our report is the first to document intercellular diversity of expression of a specific sugar linkage (α2,6-Sial) *in vivo*.

**Figure 1:**
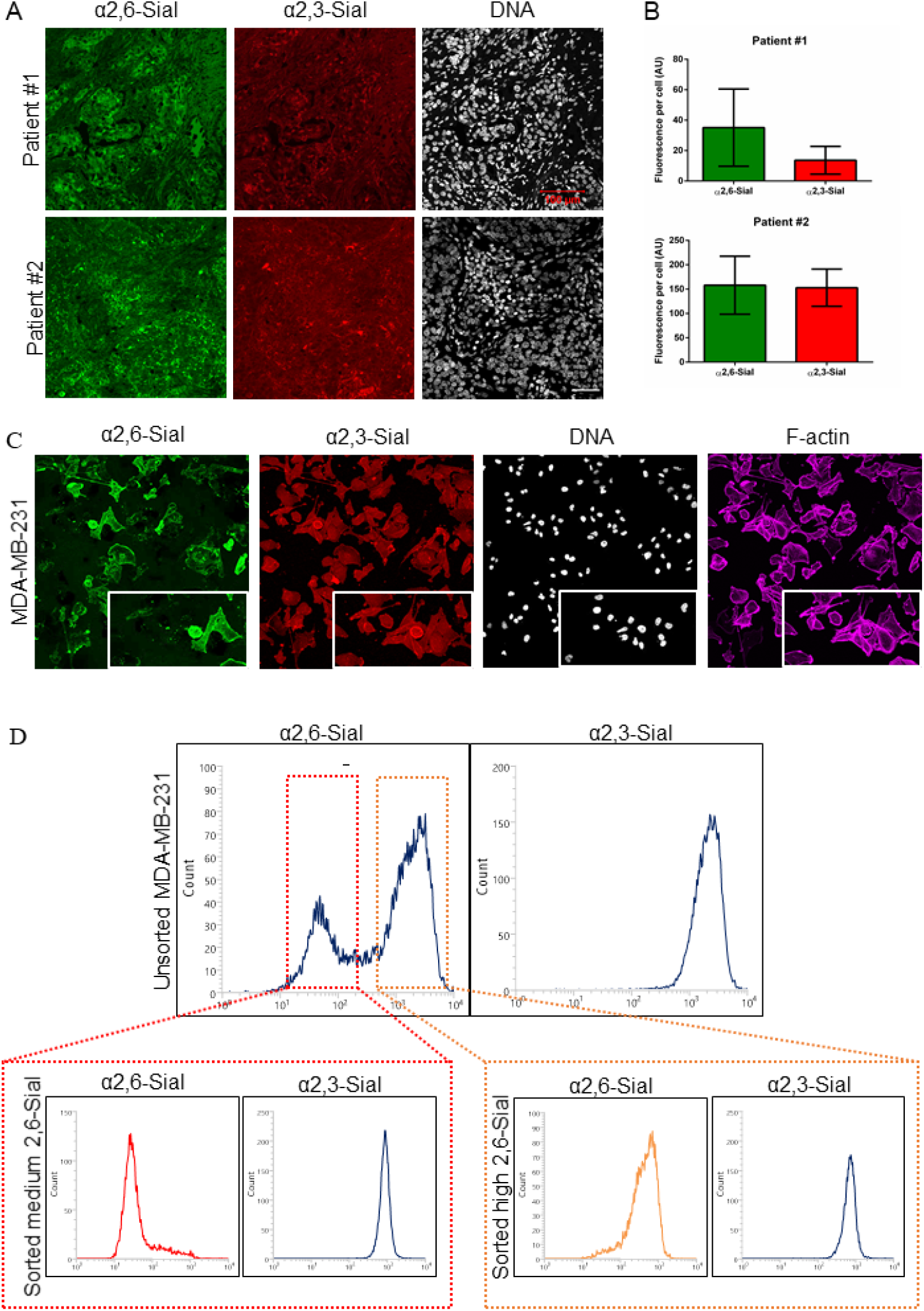
α2,6-sialic acid heterogeneity in breast cancer. (A) Confocal micrographs showing α2,6-sialic acid (SNA-FITC, green) and α2,3-sialic acid (MAA-TRITC, red) staining of breast cancer sections from two individuals (top and bottom rows) showing heterogeneity in α2,6-sialic acid linkage expression. Nucleus is stained with DAPI (white) (scale bar: 100 μm). (B) Bar graphs showing quantification of individual sialic acid levels from breast cancer sections shown in 1A. Error bars represent mean ± SD. (C) Confocal micrographs showing heterogeneity in α2,6-Sial levels (SNA-FITC green) and uniform α2,3-Sialic levels (MAA-TRITC, red) in invasive breast cancer cell line MDA-MB-231 using lectin cytochemical fluorescence. Cells are counter stained for nucleus with DAPI (white) and F-actin with Phalloidin (Magenta) Insets of a subfield within the images shown in bottom right corner (D) Lectin-based flow cytometry profiles of MDA-MB-231 cells showing bi-modal distribution of α2,6 Sial levels on (top panel, left) and α2,3 Sial levels showing uni-modal distribution (top right). Red inset shows moderate levels of α2,6-Sial (left) and unchanged α2,3-Sial (right) in sorted medium 2,6-Sial cells. Orange inset shows higher α2,6-Sial (left) and unchanged α2,3-Sial (right) levels in sorted high 2,6-Sial cells.

We next asked whether such heterogeneous expression could also be observed within cultured breast cancer cell lines. Lectin cytochemistry of the breast cancer cell line MDA-MB-231, using FITC-conjugated SNA and TRITC-conjugated MAA revealed higher signals (Fig 1C) compared to lactose sugar controls (Fig S2). Similar to our *in vivo* findings, we observed marked variation in α2,6-Sial linkage levels between MDA-MB-231 cells in the same field (Fig 1C Green. Inset). Such variations were not appreciable for α2,3-Sial levels between the same cells (Fig 1C Red, Inset). Combining lectin-binding with flow cytometry, we were able to discern two subpopulations of MDA-MB-231 cells with distinct levels of α2,6-Sial, evident from a bimodal distribution of the staining intensity histogram (unstained cells or cells stained with FITC were used as negative control (Fig S3&S4) (a third subpopulation is a minor fraction that does not stain for α2,6-Sial; Fig 1D, Left). The level of α2,3-Sial in the same population showed a sharp unimodal distribution (Fig 1D, right) confirming our observation from Figure 1C. We then sorted these subpopulations based on α2,6-Sial as shown in Figure 1D and will refer to them from hereon as low 2,6-Sial- (cells that did not stain for α 2,6-Sial), medium 2,6-Sial- and high 2,6-Sial- cells. After sorting, the individual populations were cultured separately. In early passages, the medium 2,6-Sial subpopulation continued to show a sharp and unimodal peak of moderate α2,6-Sial level coincident with the first peak of the bimodal distribution seen in unsorted MDA-MB-231 cells (Fig 1D red inset, left). Early passage high 2,6-Sial cells also showed a unimodal peak of α2,6-Sial coincident with the second peak of the bimodal distribution seen in the unsorted MDA-MB-231 cells (Fig 1D orange inset, right). To our surprise, low 2,6-Sial cells, upon culture, shifted to stably express α2,6 Sial to levels concurrent with the medium 2,6-Sial subpopulation (Fig S5 green inset, left). All the three subpopulations showed similar levels of α2,3-Sial (Fig 1D red inset, orange inset, right and Fig S5 green inset, right). We confirmed the differential levels of α2,6-Sial levels in the sorted, early passage cultures of the high- and medium-2,6-Sial cells using lectin cytochemical fluorescence (Fig S6, green), even though α2,3-Sial levels remained unchanged between them (Fig S6, red)). We also probed for other glycans in unsorted MDA-MB-231 cells. Levels of bisecting, bi- (Fig S7, green, top row), tri-, and tetra-antennary N-linked glycans (Fig S7, green, middle row) as well as Core 1, mono/di sialyl Core 1 O-linked glycans (Fig S7, green bottom row) showed uniform expression. Similarly, sharp unimodal distribution was observed when the sorted high- and medium-2,6-Sial cells were probed for the above glycans (Fig S8). These findings suggest that the intercellular heterogeneity that we observe for α2,6-Sial is terminal glycan-and linkage-specific and is not the outcome of a deeper heterogeneity in N- or O-glycan core biosynthesis.

### Medium 2,6-Sial cells show greater plasticity and invasion than high 2,6-Sial cells

We next sought to investigate the functional differences between the high- and medium- 2,6-Sial subpopulations. Upon serial passaging in culture, medium 2,6-Sial cells showed a gradual recapitulation of the bimodal expression seen in flow cytometry of unsorted MDA-MB-231 cells (Fig 2A). On the other hand, the high 2,6-Sial cells showed a consistent high and unimodal level of α2,6-Sial (Fig 2B). We did not observe any changes in α2,3-Sial levels in both medium and high 2,6-Sial cells upon serial passaging (Fig S9&S10).

**Figure 2:**
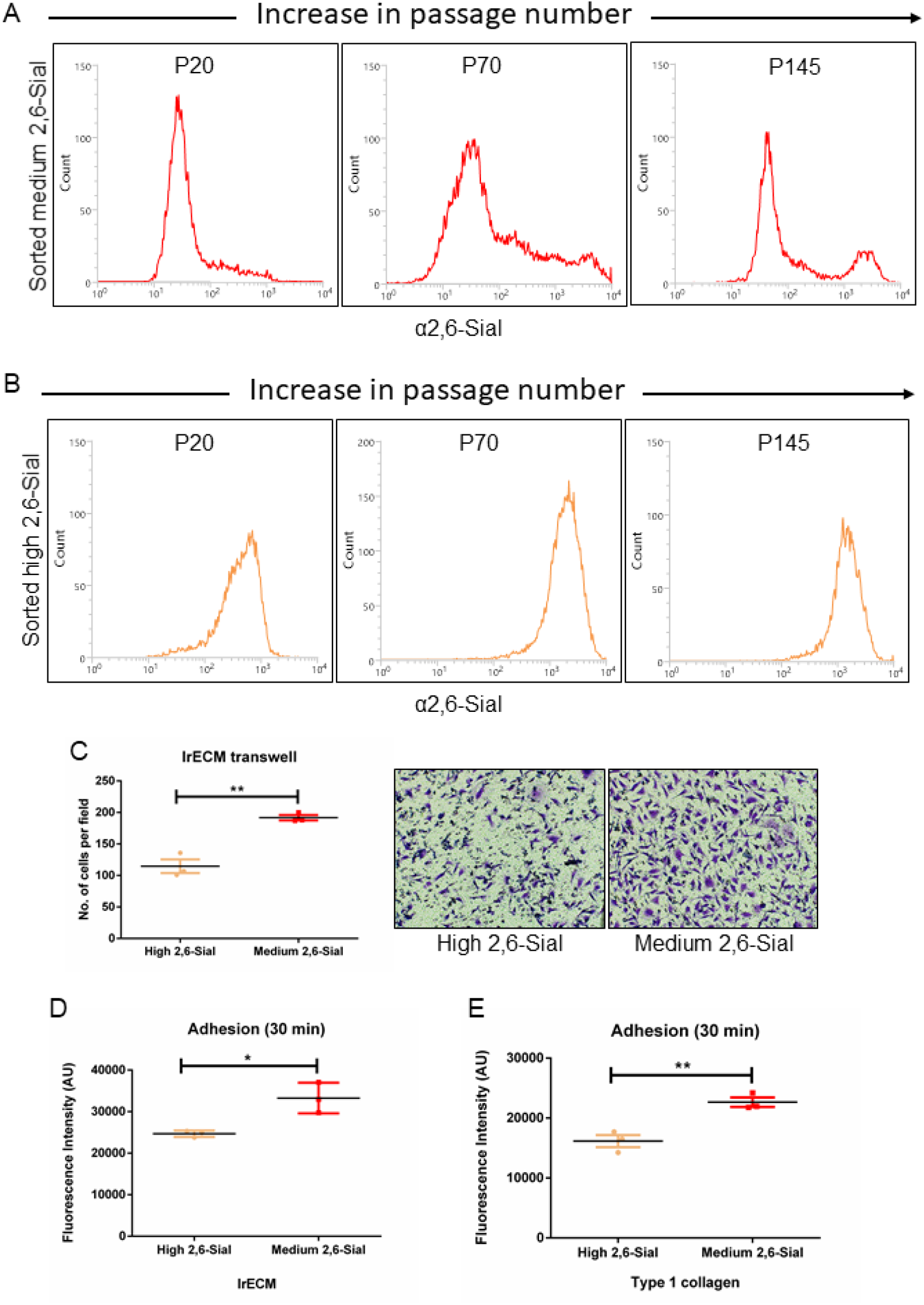
Medium- 2,6-Sial cells show greater plasticity adhere better to and invade through ECM. (A) Lectin-based flow cytometry profile of α2,6- Sial levels of medium 2,6-Sial population at three passages (20, 70 and 145) of long-term culture showing a gradual recapitulation of bimodal α2,6-Sial distribution after 70 passages. (B) Lectin-based flow cytometry of high 2,6-Sial at three passages (20, 70 and 145) in long-term culture showing no change in α2,6-Sial even after 145 passages. (C) Bar graph showing lower invasion of high 2,6-Sial cells (yellow) compared with medium 2,6-Sial cells (red) that invaded to the other side of lrECM-coated transwells. (n=3)). (D) Graph showing lower adhesion of high 2,6-Sial cells (yellow) compared with medium 2,6-Sial cells (red) to lrECM (n=3)). (E) Graph showing lower adhesion of high 2,6-Sial cells (yellow) compared with medium 2,6-Sial cells (red) to Type 1 Collagen (n=3). Error bars denote mean ± SEM. Unpaired Student’s *t* test was performed for statistical significance (\**P*<0.05).

Next, we investigated the ability of these subpopulations to invade through ECM. Medium 2,6-Sial cells invaded more through lrECM (laminin rich ECM also called as laminin rich basement membrane (lrBM)) coated transwells, when compared with high 2,6-Sial cells (Fig 2C). Invasion of mesenchymal cells requires strong tethering to the matrix substrata (30). Therefore, the adhesion of the sorted populations to both the non-fibrillar laminin rich BM matrix (lrBM) and the fibrillar Type 1 collagen was assessed. For both matrices, higher adhesion was observed for medium 2,6-Sial cells when compared to high 2,6-Sial cells. (Fig 2D and 2E; BSA-coated surface, as a negative control showing negligible cell adhesion (Fig S11)).

### Medium 2,6-Sial cells invade and disperse further than high 2,6-Sial cells

The ability of medium 2,6-Sial cells to adhere better to and invade more through ECM than high 2,6-Sial cells led us to hypothesize if the former invades in collective manner or through solitary mesenchymal movement where adhesion to ECM is crucial. To answer the question, we used a customized 3D assay, wherein clusters of cancer cells are coated with lrBM matrix and then embedded within fibrillar Type 1 collagen to mimic the collagen-rich stromal environment (31). In concurrence with our transwell experiments, single medium 2,6-Sial cells (Fig 3A) were found to radially invade into the Type 1 collagen to a greater extent than high 2,6-Sial cancer cells (Fig 3B). Time-lapse bright field microscopy on such cultures allowed us to measure the radial collective cellular migration as well as count the single cells that migrated into the fibrillar ECM (Fig 3C; Video S1A&B). Both medium- and high- 2,6-Sial cells showed comparable collective cell migration (Fig 3D). However, the number of single medium 2,6-Sial cells that invaded into the collagen as well as their mean migratory velocity was significantly higher than that of high 2,6-Sial cells (Fig 3E & F).

**Figure 3:**
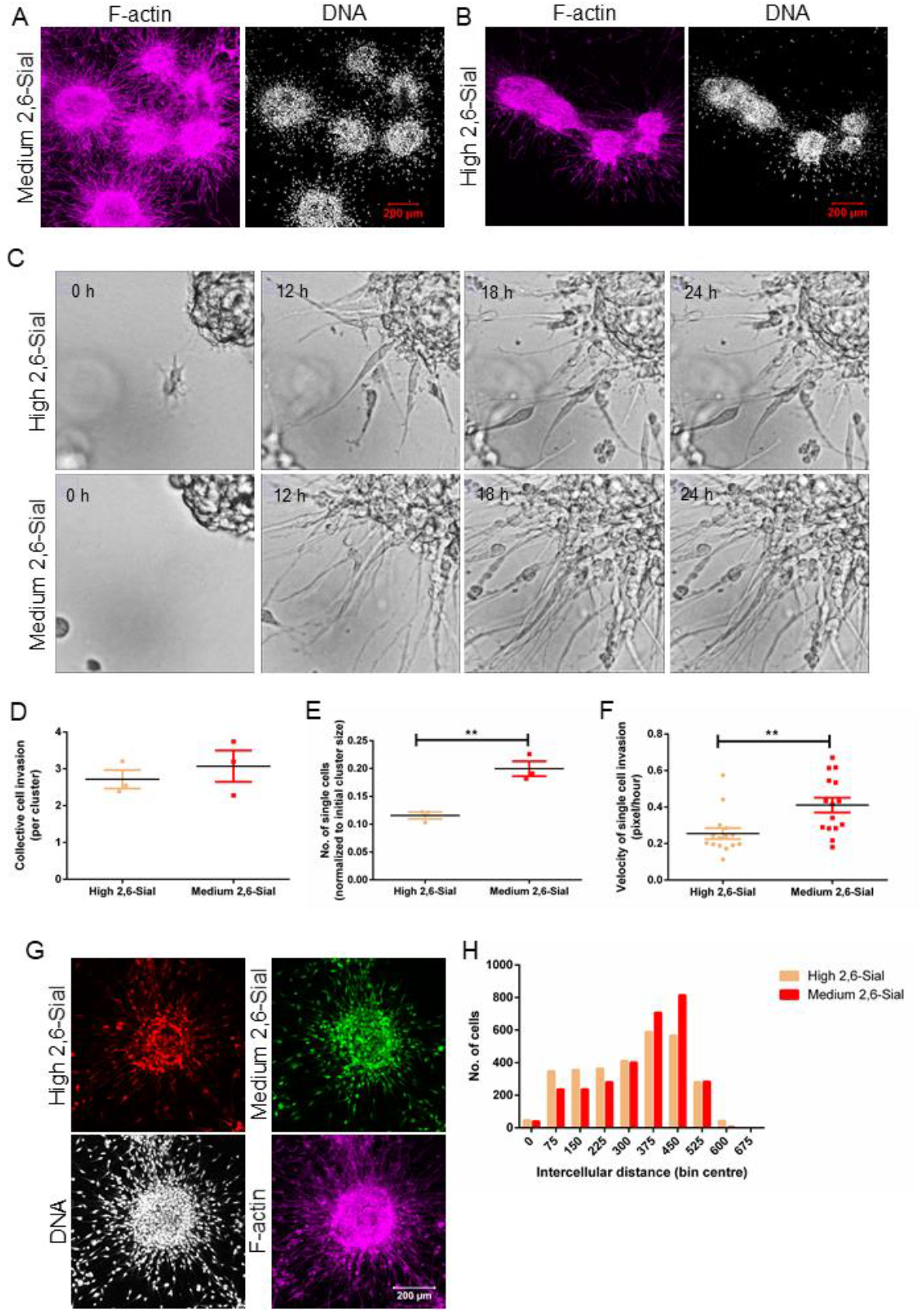
Medium 2,6-Sial cells invade faster through a 3D pathotypic multi-ECM microenvironment. (A) Confocal micrographs showing medium 2,6-Sial cells invading into fibrillar Type 1 collagen matrix from lrECM-coated multicellular clusters after 24 h (B) Confocal micrographs showing high 2,6-Sial cells invading into fibrillar Type 1 collagen matrix from lrECM-coated multicellular clusters after 24 h (A,B: Cells are counter-stained for nucleus with DAPI (white) and F-actin with Phalloidin (Magenta)(Scale bar: 200 μm). (C) Bright field images taken at 0h, 12h, 18h and 24h from time-lapse videography of lrECM-coated clusters of high- and medium- 2,6-Sial (top and bottom) invading into surrounding Type 1 collagen. (D) Graph showing insignificant differences in collective cell mode of invasion of high (yellow) and medium 2,6-Sial cells (red) as measured by increase in cluster size obtained from time lapse videography (n=3) (E) Graph showing significantly lower invasion of high 2,6-Sial cells (yellow) compared with medium 2,6-Sial cells (red) as measured by the number of dispersed single cells in Type 1 Collagen normalized to the initial cluster size obtained from lapse videography (n=3) (F) Graph showing significantly lower mean migratory velocity of single high 2,6-Sial cells (yellow) compared with medium 2,6-Sial cells (red) as measured by manual tracking dynamics obtained from lapse videography (n=3, N>=15 cells). (G) Confocal micrographs showing differential sorting of medium 2,6-Sial (green) cells invading and dispersing further into the Type 1 collagen ECM while high 2,6-Sial cells (red) form the core of the cluster when these two cells have been mixed in equal proportion and cultured in 3D. (Cells are counter-stained for nucleus with DAPI (white) and F-actin with Phalloidin (Magenta) (Scale bar: 200 μm). (H) Histogram showing distribution of intercellular distances between high 2,6-Sial cells (red) compared with medium 2,6-Sial cells (yellow). Intercellular distances between medium α2,6-Sial cells are shifted rightwards indicative of a greater spread and farther invasion within 3D matrix microenvironment compared to high 2,6-Sial cells Error bars denote mean ± SEM. Unpaired Student’s *t* test was performed for statistical significance (\**P*<0.05, \*\**P*<0.01).

This prompted us to ask whether differential invasion would cause a population with both high- and medium- 2,6-Sial cells in a cancer cluster embedded in ECM to self-sort, with the former giving rise to the indolently growing bulk and the latter forming the radially invading front. This was indeed found to be the case (Fig 3G): when the two subpopulations were labelled with constitutively expressing fluorescence reporters, mixed in equal numbers and embedded in BM and Type 1 collagen, medium 2,6-Sial cells (Fig 3G, green) predominantly were present within the collagen and high 2,6-Sial cells (Fig 3G, red) were closely clustered together in the central core. The relatively higher dispersion of the medium 2,6-Sial cells was confirmed by plotting the intercellular distances separately for medium- and high- 2,6-Sial cells wherein the histogram for the latter showed a leftward skew relative to the former (Fig 3H)

### Medium- and high- 2,6-Sial cells differ in their expression of *ST6GAL1*

We next asked whether the distinct α2,6-Sial levels in the two sorted populations could be due to differential expression of genes involved in glycan synthesis and/or sialic acid metabolism. To do so we examined the expression of genes encoding proteins involved in N-linked glycan synthesis (*ALG1*, *MAN1A1, DPAGT1, ALG3 and GANAB*), sialic acid synthesis and sialidase (*CMAS, GNE, NANS and NEU1*), and those coding for 2,3- and 2,6- sialyl transferases (Fig 4A) using quantitative real-time PCR (qRT-PCR). We did not observe any significant change in the expression of genes involved in N-linked glycosylation and sialic acid synthesis, sialidase (Fig 4B&C) confirming our flow cytometry results on detection of differences in N-glycans between the two sorted populations. The expression of genes encoding 2,3-sialyltransferases was not significantly changed confirming equivalent expression of 2,3-Sial levels in the medium- and high- 2,6-Sial cells (Fig 4C). We found that *ST6GAL1* mRNA levels were lower in medium 2,6-Sial cells when compared with high 2,6-Sial cells. Expression levels of other α2,6 sialyl transferase genes like *ST6GALNAC2*, *ST6GALNAC4*, *ST6GALNAC6* did not vary significantly between populations (Fig 4D).

**Figure 4:**
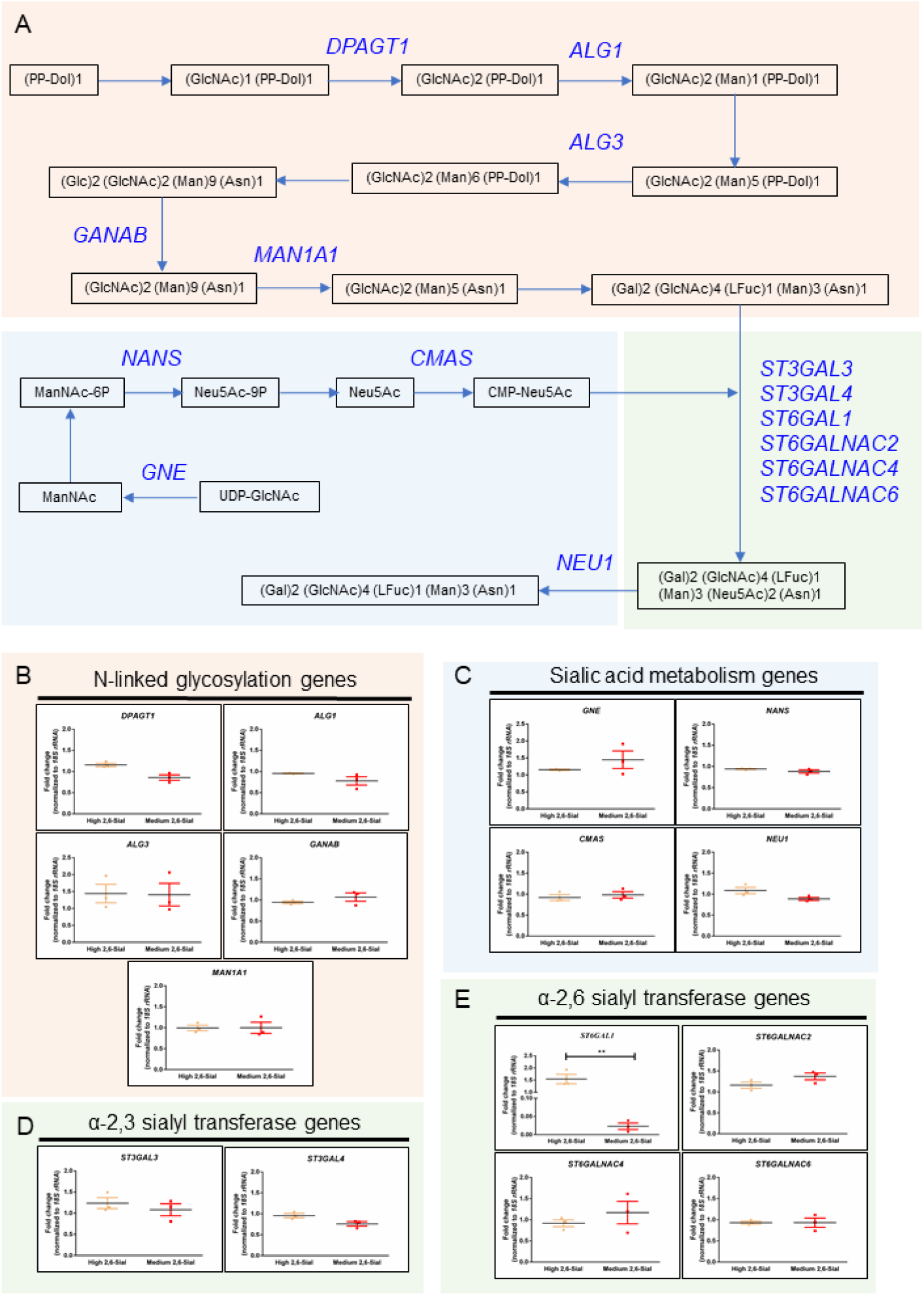
Differential expression of *ST6GAL1* gene. (A) Schematic depiction of key processes involved in the fate and utilization of sialic acids and the genes encoding the enzymes involved in these processes arranged in the temporal order of their function: N-glycan synthesis, sialic acid metabolism and sialyltransferases. (B) Graphs depicting relative mRNA levels of genes involved in N-linked glycosylation *DPAGT1*, *ALG1*, *ALG3*, *GANAB*, *MAN1A1*; expression is insignificantly altered between high (yellow) & medium 2,6-Sial cells (red). (C) Graphs depicting relative mRNA levels of genes involved in sialic acid metabolism *GNE*, *NANS*, *CMAS*, *NEU1*; expression is insignificantly altered between high (yellow) & medium 2,6-Sial cells (red). (D) Graphs depicting relative mRNA levels of genes involved in 2,3-Sialic acid conjugation *ST3GAL3* and *ST3GAL4*; expression is insignificantly altered between high (yellow) & medium 2,6-Sial cells (red). (E) Graphs depicting relative mRNA levels of genes involved in sialic acid metabolism *ST6GAL1*, *ST6GALNAC2*, *ST6GALNAC4* and *ST6GALNAC6*; expression is insignificantly altered between high (yellow) & medium 2,6-Sial cells (red) for all except *ST6GAL1*, which is significantly lower in medium 2,6-Sial cells compared to high 2,6-Sial cells. Expression of all genes is expressed as fold change normalized to unsorted cells. *18S rRNA* gene is used as internal control. Data shown is from three independent biological experiments with at least duplicate samples run in each experiment. Error bars denote mean ± SEM. Unpaired Student’s *t* test was performed for statistical significance (\*\**P*<0.01).

### Knockdown of *ST6GAL1* in high 2,6 Sial cells enhances their mesenchymal invasion

To establish if α2,6 sialic acid linkage has a direct effect on cancer cell invasion, we knocked down the expression of *ST6GAL1* in high 2,6-Sial cells using lentivirally delivered shRNA. *ST6GAL1* expression was confirmed using qRT-PCR (Fig S12). Lectin flow cytometry confirmed a resultant decrease of α2,6-Sial levels in high 2,6-Sial cells with *ST6GAL1* knockdown (sh*ST6GAL1*-high 2,6-Sial cells) (Fig 5A, left) compared with scrambled control cells (shSc-high 2,6-Sial cells) (Fig 5A, middle). As a result of the knockdown, α2,6-Sial levels of sh*ST6GAL1*-high 2,6-Sial cells were comparable to medium 2,6-Sial cells (Fig 5A, left & right).

**Figure 5:**
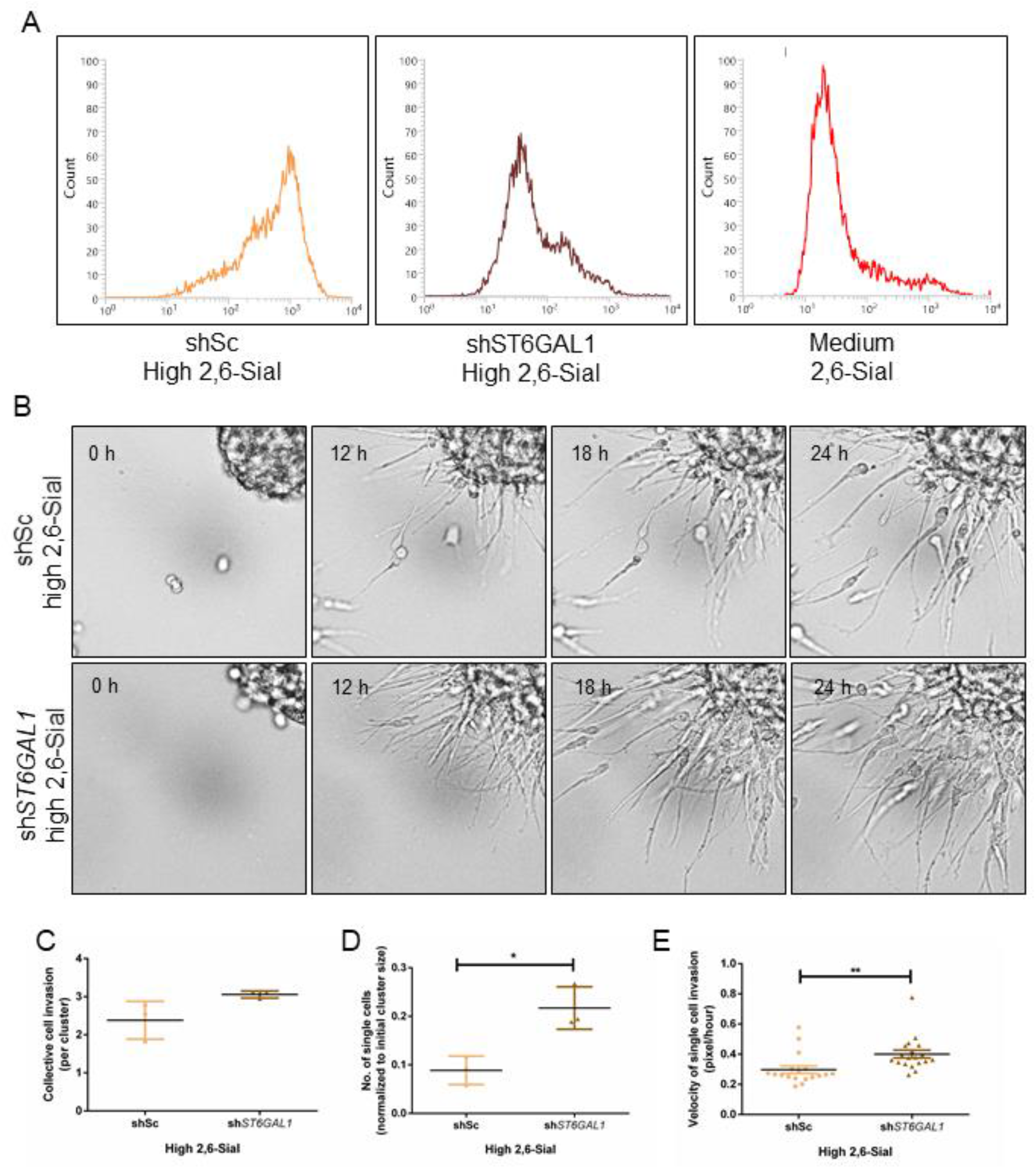
α2,6-Sial levels regulate mesenchymal invasion of high 2,6-Sial cells. (A) Lectin-based flow cytometry profiles showing decreased α2,6-Sial levels upon *ST6GAL1* gene knockdown in high 2,6-Sial cells. (B) Bright field images taken at 0h, 12h, 18h and 24h from time-lapse videography of lrECM-coated clusters of scrambled control- and *ST6GAL1* knocked down-high 2,6-Sial cells (shSc high 2,6-Sial top and sh*ST6GAL1* high 2,6-Sial bottom) invading into surrounding Type 1 collagen. (C) Graph showing insignificant differences in collective cell mode of invasion of shSc high 2,6-Sial (yellow) and sh*ST6GAL1* high 2,6-Sial cells (brown) as measured by increase in cluster size obtained from time lapse videography (n=3) (D) Graph showing significantly lower invasion of shSc high 2,6-Sial cells (yellow) compared with sh*ST6GAL1* high 2,6-Sial cells (red) as measured by the number of dispersed single cells in Type 1 Collagen normalized to the initial cluster size obtained from lapse videography (n=3) (E) Graph showing significantly lower mean migratory velocity of single shSc high 2,6-Sial cells (yellow) compared with sh*ST6GAL1* high 2,6-Sial cells (red) as measured by manual tracking dynamics obtained from lapse videography (n=3, N>=15 cells). Error bars denote mean ± SEM. Unpaired Student’s *t* test was performed for statistical significance (\**P*<0.05, \*\**P*<0.01).

When assayed for invasion from within lrBM to Type 1 collagen, sh*ST6GAL1*-high 2,6-Sial cells showed greater presence in collagen compared to shSc-high 2,6-Sial cells (Video S2A and 2B) (Fig 5B). Collective cell invasion did not show any difference between knockdown and scrambled control cells (Fig 5C). However, the number of invaded single mesenchymal sh*ST6GAL1*-high 2,6-Sial cells in the collagen and their mean migratory velocity in ECM was greater than shSc-high 2,6-Sial cells (Fig 5D&E). These observations suggest that single cell invasion of breast cancer epithelia may be directly regulated by their surface α2,6-Sial levels.

### A computational model shows that altered matrix-adhesion dynamics is sufficient to explain differential invasion of cancer epithelia

Is greater adhesion of medium 2,6-Sial cancer cells to ECM causal to its enhanced ability for invasion when cultured in 3D separately, or in coculture with, high 2,6-Sial cells? To answer this, we resorted to a recently constructed computational model of cancer cell invasion (31) using the Compucell 3D simulation framework (32). Our model was demonstrated to simulate single cell- and collective-cell migration (individually and in combination, known as multiscale invasion) first through lrBM-like barriers and then collagen-like fibrillar environments, similar to the 3D invasion assay used above. The cellular constituents of our model were digital medium and high 2,6-Sial cells (mimicking medium- and high 2,6-Sial cells) with the former differing from the latter in exhibiting greater matrix adhesion. The ECM constituents of the model were digital lrBM and digital collagen (mimicking BM and Type 1 collagen) with the only difference being their nonfibrillar and fibrillar structure, respectively. In consonance with our experiments, digital medium 2,6-Sial cells showed greater invasion through model ECM than digital high 2,6-Sial cells (Fig 6A and 6B).

**Figure 6:**
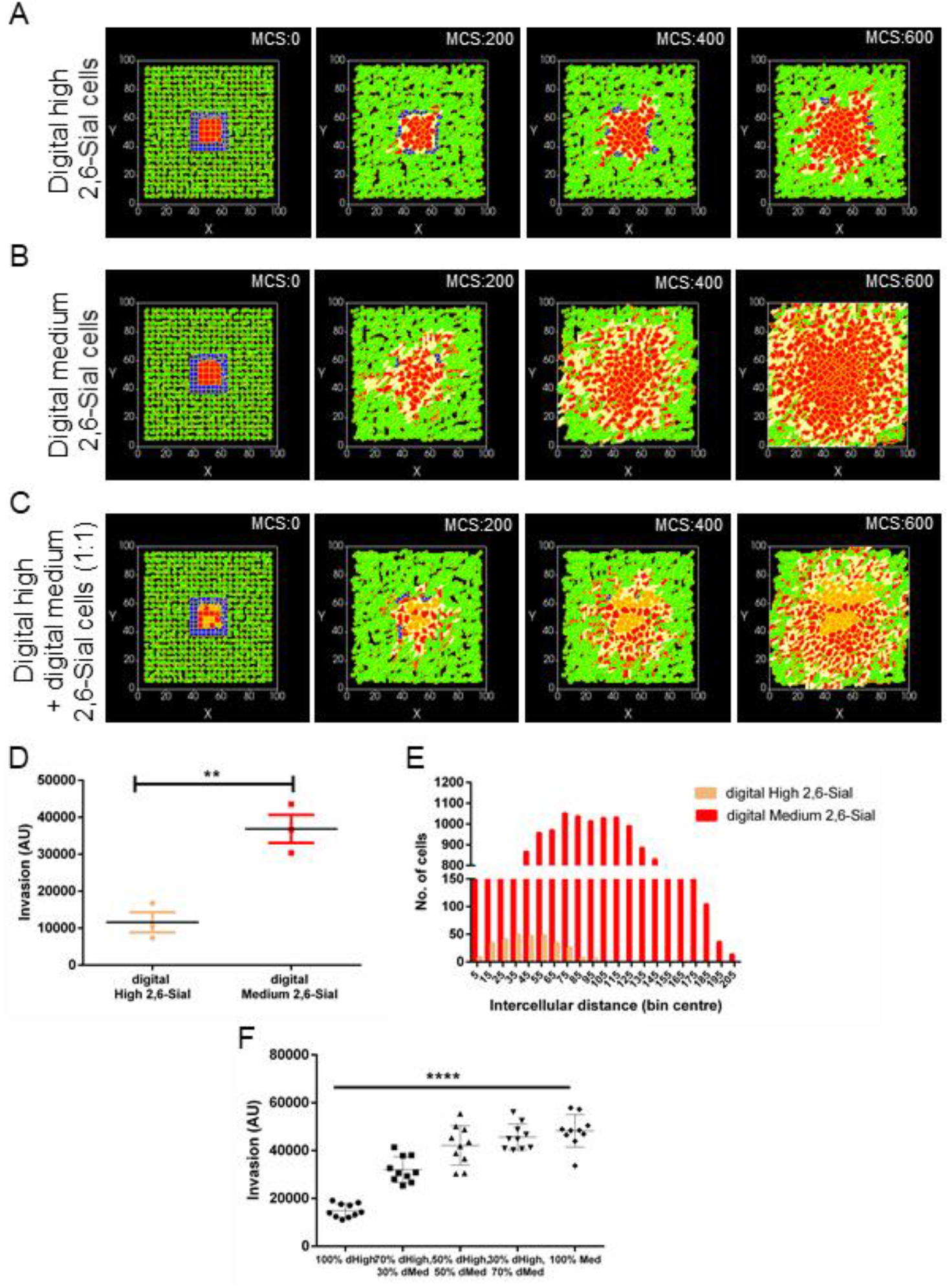
Multiscale simulations predict matrix adhesion principally contributes to increased invasion. (A) Snapshots of simulation at MCS (= 0, 200, 400 and 600) of model lrECM (blue) coated clusters of digital high 2,6-Sial cells in model Type 1 Collagen (green) (B) Snapshots of simulation at MCS (= 0, 200, 400 and 600) of model lrECM (blue) coated clusters of digital medium 2,6-Sial cells in model Type 1 Collagen (green) (C) Snapshots of simulation at MCS (= 0, 200, 400 and600) of model lrECM (blue) coated clusters of digital high and medium 2,6-Sial cells mixed in a ratio of 1:1 show a relatively greater invasion and dispersal of digital medium 2,6-Sial cells with digital high 2,6-Sial cells forming the central core. (D) Bar graph showing greater invasion digital medium 2,6-Sial cells (red) compared with digital high 2,6-Sial cells in their 3D cocultures such as in Figure 6C (n=3) Error bars denote mean ± SEM. Unpaired Student’s *t* test was performed for statistical significance. (E) Histograms depicting the intercellular distances of the digital high 2,6-Sial cells (yellow) and that of medium 2,6-Sial cells (red) with the rightward shift of the latter indicating greater dispersal. (F) Graph depicting the invasion of an overall population of cancer cells in 3D (conditions similar to Figure 6A-C) wherein the clusters of digital cells are comprised of digital high- and medium- 2,6-Sial cells in the relative proportion of 100%,0%; 75%,25%; 50%,50%; 25%,75%; and 0%,100%, from left to right respectively. Error bars denote mean ± SEM. Ordinary one-way ANOVA with Tukey’s posthoc multiple comparisons was performed for statistical significance (\*\*\*\**P*<0.0001).

We next asked whether the computational model would predict the differential radial invasion that the medium 2,6-Sial cells showed, when cultured in an admixture with high 2,6-Sial cells. Even in our simulation framework, digital medium 2,6-Sial cells differentially migrated further to form the invasive front of mixed digital tumoroid populations, wherein digital high 2,6-Sial cells formed the core (Fig 6C; quantification of individual digital medium- and high- 2,6-Sial cell invasion when present within the initial mass in a ratio of 1:1 shown in Fig 6D and the further dispersion of digital medium 2,6-Sial cells confirmed through intercellular distance histogram shown in Fig 6E). Given that the digital cells differ only with respect to their adhesion to the model ECM, the latter property is likely the determinant factor for the differential and greater invasion of real medium 2,6-Sial cells. Upon varying the ratio of digital medium- and high- 2,6-Sial cells constituting similar-sized initial clusters in our model, we observed that overall invasion was lowest in starting tumor clusters with 100% digital high 2,6- Sial cells (Fig 6F). Clusters with 70% digital high- and 30% digital medium 2,6 Sial cells invaded to a greater extent than 100% high clusters. Clusters with 50% high and medium, 30% high and 70% medium, and 100% medium 2,6-Sial cells invaded higher than both the previous two conditions but showed no significant difference with respect to each other. The distribution of invasion across 10 runs for these three conditions showed that the arrangement of cells within such clusters and the stochasticity of multiple interactions between cells and ECM during the simulation run contributed to the magnitude of the phenotype. Notably, there were certain arrangements where clusters with as less as 50% digital medium cells ‘out-invaded’ 100% digital medium 2,6-Sial cell clusters.

## Discussion

In this current study, we establish for the first time the occurrence of intercellular heterogeneity in the expression of a specific sialic acid linkage within malignant tumors and cancer cell lines of breast. We show that cellular populations with distinct α2,6-Sial levels coexist together within cell lines and may reflect linkage heterogeneity within tumor cell populations *in vivo*. The bimodal expression of α2,6-Sial shown here can also be evidenced in several reports for MDA_MB-231 as well as other cell lines but has not been investigated before (33–37). The α2,6-Sial heterogeneity is counterpoised with relatively homogeneous expression in the same cells for other glycans such as α2,3-Sial, fucose, T/Tn- antigen, bisecting/ biantennary complex N-glycans & tri/tetra antennary complex N-glycans. This suggests that glycan heterogeneity within tumor populations may be specific to identity of monosaccharides and their linkage to preceding glycan moieties.

Several published reports note hypersialylation of cancer cells, including that of the neoplasms of breast (22) cancer. Hypersialylation is associated with, and suggested to be causal to, increased aggressiveness, stemness, resistance to chemotherapeutic agents, ability to survive in stressful conditions like hypoxia and impaired nutrient supply (24, 25, 27, 38). However, when we performed flow cytometry to isolate the two subpopulations showing distinct α2,6 sialic acid levels, the one showing lower levels (which we denoted as medium 2,6-Sial cells) showed greater invasion than high 2,6-Sial cells in both transwell assay and in 3D cultures. Medium 2,6-Sial cells were also observed to adhere better to both laminin-rich and collagenous ECM as well as migrated through the latter with higher velocity compared to high 2,6-Sial cells. When cultured together, medium 2,6-Sial cells migrated farther, and were dispersed to a greater extent than high 2,6-Sial cells. When α2,6-Sialic acid levels in high cells were genetically perturbed to levels comparable with medium 2,6-Sial cells, their velocity and dispersal in ECM increased. Can the difference in invasion be explained by an appropriate difference in adhesion to ECM?

To answer this question, we used Cellular Pott’s model-based computational simulations, which predicted that cells with higher adhesion to ECM are able to invade better into a surrounding stroma-like fibrillar environment. Our simulations also were able to provide an answer to another question: what advantage do high 2,6-Sial cells confer to tumors in the presence of the more invasive medium 2,6-Sial cells? Simulations performed with different beginning ratios of digital medium- and high- 2,6-Sial cells showed that the presence of a population of slow invading high 2,6-Sial cells could bias the invasion of medium 2,6-Sial cells in an outward direction. On instances, such mixed cell populations could invade more than populations solely consisting of medium 2,6-Sial cells. The inertial behaviour exhibited by high 2,6-Sial cells is prognostic of jamming-unjamming dynamics proposed to play an important role in the physical mechanisms of cancer progression and suggests that the sialic acid heterogeneity could give rise to heterogeneity in material behaviour of tumor subpopulations. We will actively investigate this aspect in the future (39).

Our observations made through the assessment of endogenous expression of *ST6GAL1* (which is positively correlated with 2,6-Sial levels), serves to reconcile the contradiction between reports of increased invasion as a result of forced overexpression of *ST6GAL1* within cancer cells (24, 27) and an overall decreased level of *ST6GAL1* expression in breast cancer tissues assessed within The Cancer Genome Atlas (40). We posit that within a population with heterogeneous expression of α2,6-sialic acids, those with a moderate expression will escape faster through a mesenchymal invasive process. At the same time, the medium 2,6-Sial cells will continue to give rise to the high 2,6-Sial populations, as demonstrated by our flow cytometric experiments on continuously passaged sorted cells. In conclusion, our results show that the intercellular sialic acid heterogeneity and breast cancer cell invasion are not just spatiotemporally coincident, these two processes actively drive each other. It will be imperative to break the link between the two by targeting sialic acids through novel therapeutic strategies such as precision glyco-editing (41) and sialic acid-siglec-based immunotherapy (42, 43).

## Materials and Methods

### Cell culture

MDA-MB-231 cells were maintained in DMEM:F12 (1:1) (HiMedia AT140) along with 10% fetal bovine serum (Gibco, 10270) in a 5% carbon dioxide, 37°C temperature humidified incubator.

### Lectin histochemistry

Breast tumor and normal sections were made from paraffin embedded blocks ay Kidwai cancer institute, Bangalore after obtaining necessary approval from Institutional Human Ethics committee and consent from patients. Sections were incubated at 65 °C overnight to remove wax. Immediately, samples were re-hydrated gradually incubating in decreasing concentrations of alcohol: 2x 5 min Xylene, 2x 5 min 100% Ethanol, 2x 5 min 90% Ethanol, 1x 10min 80% Ethanol, 1x 10 min 70% Ethanol and finally in distilled water for 10 min. Antigen retrieval was performed using citrate buffer pH 6.0 in microwave for 30 min and allowed to cool down to room temperature. Sections were blocked using 1X Carbo-Free™ blocking buffer (Vector labs, SP-5040) made in PBS pH 7.4 for 1 h at room temperature. Fluorescently labelled SNA (Vector Labs, FL-1301) and MAA (bioWORLD, 21511106-1) was added to sections at 1:100 dilution and incubated overnight at 4 °C. Lectins preincubated with 250 mM lactose (HiMedia, RM565G) was used as a negative control. Sections were washed with 1X PBS 5 min at room temperature thrice. Counterstain sections with 1 μg/mL DAPI (Thermo Fischer Scientific, D1306) for 10 min, wash excess stain and mount for imaging.

### Lectin cytochemistry

15,000 cells were seeded in 8-well chamber cover glass (Eppendorf, 0030742036). After 24 h, remove spent medium, wash cells with cold 1X PBS once and fix using ice cold 4% formaldehyde at 4 °C for 20 min. Remove excess fixative and incubate with 2% glycine for 30 min at room temperature to neutralize trace fixative. Wash thrice with 1X PBS and block using 1X Carbo-Free™ blocking buffer for 1 h at room temperature. Add fluorescent conjugated SNA and MAA at 1:500 dilution and incubate for 3 h at room temperature or overnight at 4 °C. Wash cells with 1X PBS for 5 min thrice. Counter stain cells with 1 μg/mL DAPI and 1:500 Alexa633 conjugated Phalloidin (Thermo Fischer Scientific, A22284) for 1 h at room temperature. Wash with 1X PBS 5 min twice and image.

### Lectin flow cytometry and sorting

MDA-MB-231 cells were trypsinized and counted. 0.3×10^6^ cells for analysis and 10^6^ cells for sorting were taken in a polypropylene FACS tube in 100 μL and 500 μL for analysis and sorting respectively in 1X Carbo-Free™ blocking buffer. Cells were incubated with fluorescently labelled SNA and MAA at 20 μg per 10^6^ cells concentration for 20 min at room temperature. Lectin incubated with 250 mM lactose overnight at 4 °C overnight or FITC was used as negative control. Finally, cells were diluted to 10^6^/ mL using Carbo-Free™ blocking buffer and analysed/sorted using BD Influx™ flow cytometer. For analysis, atleast 10,000 total events were acquired. For sorting, single cell purity mode was used. Lectins used in this study are listed in Table S1.

### ECM coating for adhesion assay

96 well plates were coated with 50 μg/mL reconstituted basement membrane (rBM or lr-BM) (Corning, 354230) or 50 μg/mL neutralized rat tail collagen (rich in type 1 collagen) (Gibco, A1048301) for 2 hours at 37 °C. Excess matrix was removed, allowed to dry for 30 min at 37 °C and blocked with 0.5% BSA (HiMedia, MB083) overnight at 37 °C. After overnight blocking, excess BSA was removed and plates are used for adhesion assay. 0.5% BSA overnight coating at 37 °C was used as a negative control.

### Adhesion assay

MDA-MB-231 high and medium α-2,6 sialic acid cells were trypsinized. After counting, 30,000 cells per well were incubated in BSA and ECM coated wells for 30 min at 37 °C. Unadhered cells were removed carefully and wells were washed with 1X phosphate buffered saline pH 7.4 (PBS) thrice to remove unadhered cells. Cells were fixed using 100% methanol for 10 min at room temperature. After fixing, cells were washed with 1X PBS thrice and stained with 50 μg/mL propidium iodide (HiMedia, TC252) for 30 min at room temperature. Remove excess stain and wash cells thrice with 1x PBS. Using plate reader, fluorescence was read using Ex 535nm/Em 617nm. BSA or ECM without cells was used as blank. Assay was done in triplicates and repeated three times independently.

### Quantitative real time PCR

Total RNA was isolated from high and medium α-2,6 Sial cells using TRIzol™ as per manufacturer’s protocol (Invitrogen, 15596078). Total RNA was quantified using UV-visible spectrophotometer (NanoDrop™, Thermo Fischer Scientific). 1 μg of total RNA was reverse transcribed using Verso™ cDNA synthesis kit as per manufacturer’s protocol (Thermo Scientific, AB-1453). Real time PCR was performed with 1:2 diluted cDNA using SYBR green detection system (Thermo Fischer Scientific, F415L) and Rotorgene Q (Qiagen, 9001560). *18 S rRNA* gene was used as internal control for normalization. Relative gene expression was calculated using comparative Ct method and gene expression was normalized to unsorted cells. All the genes analysed along with sequence is mentioned in Table S2. (I will check for sequence). Appropriate no template and no-RT control were included in each experiment. All the samples were analysed in duplicates/triplicates and repeated three times independently.

### Genetic perturbation of *ST6GAL1* gene

*ST6GAL1* gene shRNA clone was obtained from MISSION shRNA library (Sigma Merck, USA). Plasmid containing shRNA or scrambled control was packaged into lenti virus using packaging vectors pMD2.G and psPAX2 (packaging vectors were a kind gift from Dr. Deepak K Saini). The plasmids were transfected into 293FT cells (Thermo Fischer Scientific, R70007) using TurboFect™ (Thermo Fisher Scientific, R0533). Cells were cultured in DMEM supplemented with 10% FBS, conditioned medium containing viral particles was collected at 48 h and 72 h. After filtering through 0.45 μm filter, viral particles were concentrated using Lenti-X™ concentrator as mentioned in manufacturer’s protocol (TaKaRa, 631232). Concentrated virus was aliquoted and stored at −80 °C until use. High α2,6-Sial cells were seeded in a 24 well plate at 50-60% confluence and transduced with viral particles containing shRNA or scrambled control along with polybrene (4 μg/mL) for 24 h. After 72 h transduced cells were selected using 5 μg/mL puromycin (HiMedia, CMS8861). Knockdown of gene was assayed using real time PCR and lectin flow cytometry as described above.

### 3D Invasion assay

3D invasion assay was performed as described previously by our group(31). Briefly, cancer clusters were made using 30,000 cells in a polyHEMA (Sigma, P3932) coated 96 well plated, defined medium (Blaschke etal, 1994) supplemented with 4% rBM. After 48h, clusters were collected and embedded in polymerizing rat tail collagen in a chambered cover glass. 3D cultures were grown for 24 h in 37 °C humidified incubator with 5% carbon dioxide. End point imaging was done after fixing 3D cancer clusters, counter stained with DAPI, Alexa633-Phalloidin and imaged using Carl Zeiss LSM880 confocal microscope with system optimized settings. Brightfield time lapse imaging of invading cancer clusters was performed on Olympus IX73 fluorescence microscope fitted with stage top incubator (forgot name) and 5% carbon dioxide. Images were collected for 24h with every 10 min interval.

### Transwell Invasion assay

8 μm pore-size polycarbonate transwell inserts was obtained from HiMedia (TCP083). Transwells were coated with 200 μg/mL reconstituted basement membrane (rBM or lr-BM) (Corning, 354230) as per manufacturer’s protocol. In each transwell, 3 × 10^4^ cells were seeded in 200 μL of serum free DMEM:F12 (1:1) medium. Bottom well was filled with 1 mL of 10% serum containing DMEM:F12 (1:1) and incubated for 24 h at 37 °C, 5% carbon dioxide containing humidified chamber. Carefully, medium from transwell was removed, washed with 1X PBS once and cells were fixed using 100% methanol for 10 min. After, fixing, cells were washed once again with 1X PBS and non-invading cells were carefully removed with moistened cotton swab. Transwells are stained with 1% crystal violet for 15 min and washed to remove excess dye. Membranes were dried and imaged under microscope using 40X total magnification. At least 5 independent fields were imaged per transwell and number of cells were counted. Each experiment has been performed in duplicates and repeated three times.

### Compucell 3D Model and MATLAB analysis

The simulations performed were based from an earlier-established computational model in Compucell3D (31). Compucell3D (CC3D) is an environment that comprises the lattice-based GGH model(GGH: Glazier–Graner–Hogeweg also known as Cellular Potts model (CPM)) with solvers for partial differential equations in order to simulate biological processes, wherein molecules, cells and cellular ensembles may behave concurrently across distinct spatial scales (32, 44). The software divides the whole simulation lattice into ‘cells’ (collection of pixels). A specific ‘cell type’ is assigned to each of them. All cells having the same cell type share same properties. Hence, the minimalistic target tissue-system can be broken down into cell types representing the main constituents of the system. Interaction parameters between cell types can be made to approximate biological constraints between components, similar to that of the original in vitro or in vivo biological system. Such constraints or parameters regulate the simulation through the effective energy or Hamiltonian (H) calculated at each Monte Carlo Step (MCS). The initial conditions and the behavior of cells in the model have been calibrated based on the invasive behavior of cancer cells in different types of ECM and upon treatment with pharmacological agents. The chief constituents of the model include cancer cells, non-fibrillar BM matrix, fibrillar Type 1 collagen matrix, degraded and newly synthesized cancer ECM, diffusible activator of ECM (such as MMPs) and its diffusible inhibitor (TIMP) (31). Contact energies related to adhesion between each of these constituents and diffusion and cooperative interaction between the molecules determines the behavior of the simulation and the end stage phenotype.

Key differences with the previously mentioned version of the model are-

1. Simulated Cancer cells have two cell types assigned to them. They correspond to high and medium 2,6-Sial cells in the heterogeneous population.
2. The two different cancer cells have differential adhesion to ECM, where medium 2,6-Sial cells have relatively higher adhesion to all ECM cells like BM (Basement membrane, blob-like), Type 1 Collagen (collagen 1, fibrillar), newly synthesised collagen-like ECM and degraded ECM. All other biological cancer cell-like properties are same for the digital high and medium 2,6-Sial cells.

At initial spatial configuration (marked by MCS 0), both digital high and medium 2,6-SIal cells are located centrally encapsulated by BM cells and that is further surrounded by Type 1 collagen cells. After the Monte Carlo Step (MCS) reaches 600, the cancer cells already invade the matrix with reconstitution of localised matrix through reaction-diffusion dynamics of MMPs and their inhibitors such as TIMPs. Image of the simulation at MCS 600 is collected for analysis in MATLAB.

Two different methods of analysis were used; the first one ‘Area of minimal enclosing circle’, detailed in the previous publication ((31) calculates the smallest possible circle, which encloses all cells of a certain cell type and can be considered as a quantification of invasion corresponding to that cell type in that initial configuration. The collected simulation images can be binarized for digital high 2,6-Sial cells (coloured orange) or digital medium 2,6-Sial cells (coloured red) or both (orange and red; Fig 6F) depending on the analysis. The second method calculates distances between cells in simulation images binarized with respect to either high or medium cell type. After identifying centroids of the cells in the binarized image, a MATLAB function-‘pdist’ was used to calculate distances between all the pairs of centroids and plotted in histogram. (Fig 6E) [https://in.mathworks.com/help/stats/pdist.html].

### Statistical analysis

All experiments were performed in duplicates or more. All experiments are repeated thrice independently. Prism software (GraphPad Prism 6.0) was used for the generation of graphs and analysis. For all experiments, results are represented as mean ± SEM unless mentioned. For statistical analysis, Unpaired Student’s *t* test or ordinary one-way ANOVA followed by post-hoc Tukey test for comparison of multiple groups was performed. Significance (p value) is represented as *, where *≤0.05, **≤0.01, ***≤0.001, and ****≤0.001.

## Acknowledgements

We thank the Bio-imaging facility at Division of Biological Science, Indian Institute of Science for help with confocal microscopy. We thank Bhaskar Kumawat for help with biostatistics. DP is supported by Senior Research Fellowship (SRF) from MHRD, India. This work was supported by funds from the Department of Biotechnology, India [BT/PR26526/GET/119/92/2017], SERB[ECR/2015/000280] and the Wellcome Trust/DBT India Alliance Fellowship/Grant [IA/I/17/2/503312] awarded to RB. DP (Pramanik) acknowledges KVPY for the scholarship

---

**Figure S1:**
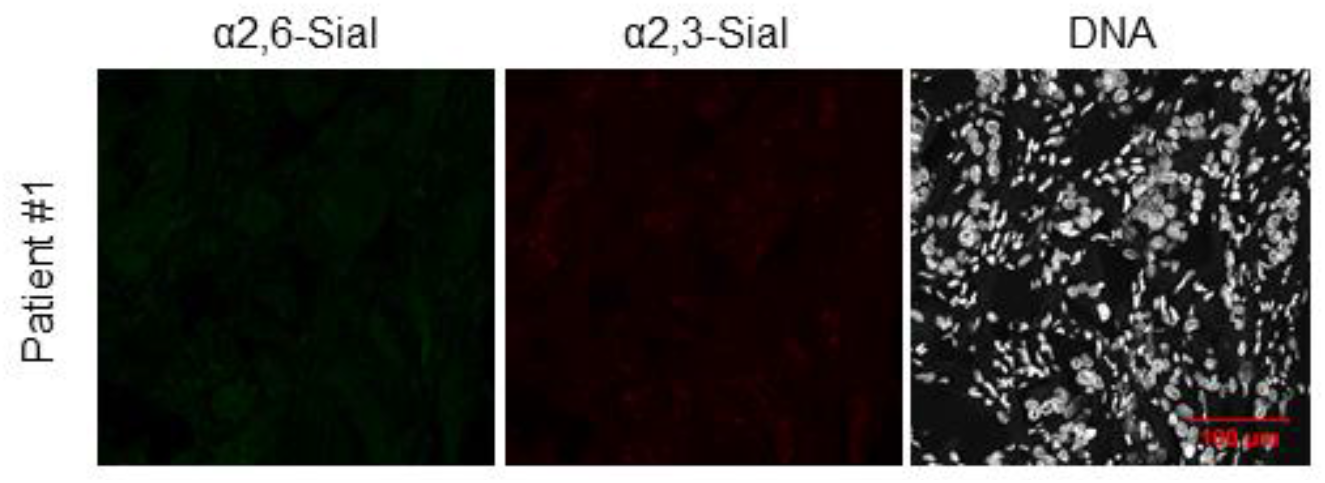
Confocal micrographs showing negligible α2,6-Sial (green) and α2,3-Sial (red) levels on breast cancer section upon staining with lectins (SNA-FITC & MAA-TRITC) pre-incubated with 250 mM lactose sugar. Nucleus is stained with DAPI (white). Scale bar: 100 μm.

**Figure S2:**
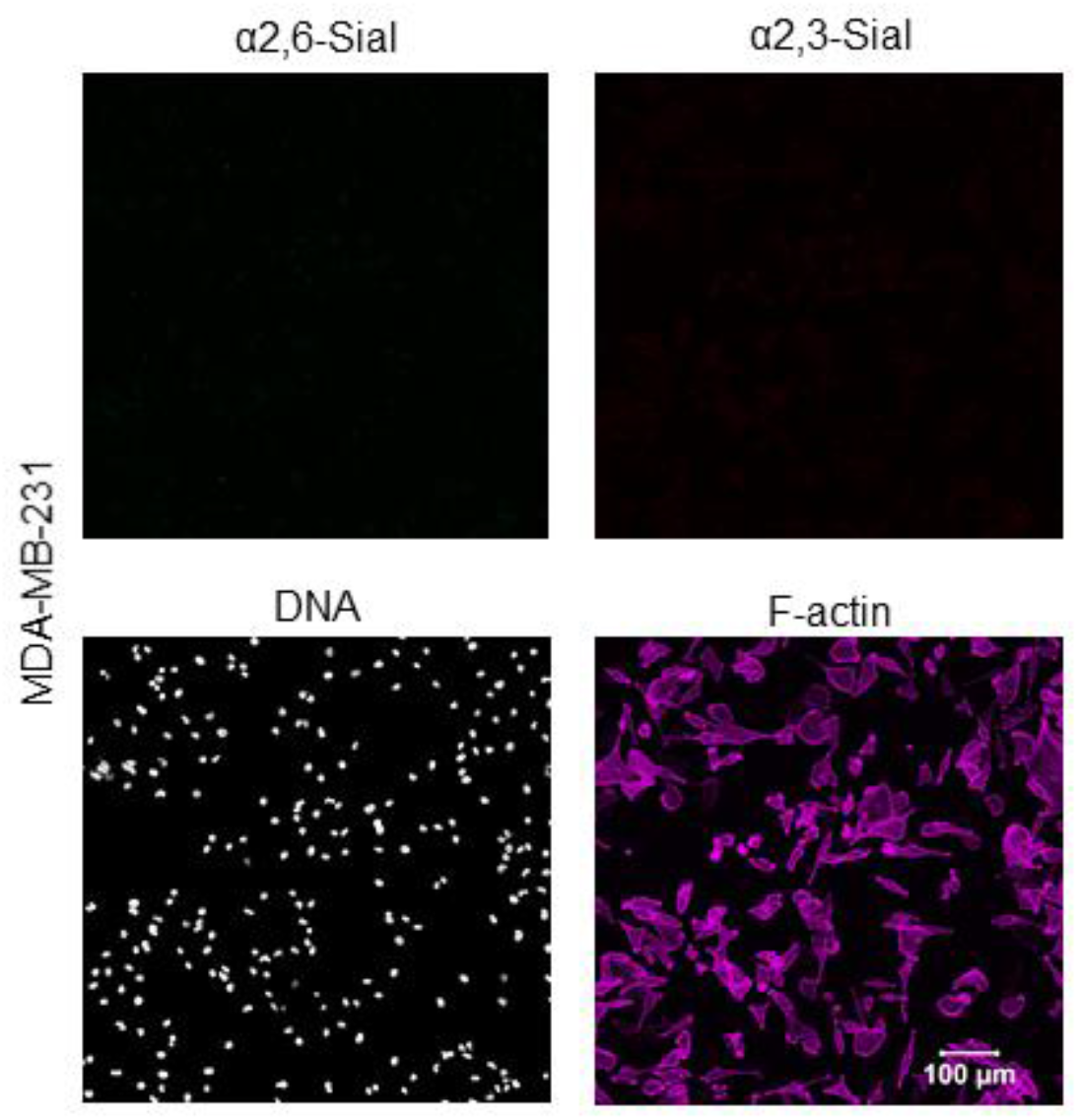
Confocal micrographs of MDA-MB-231 stained with SNA-FITC and MAA-TRITC pre-incubated with 250 mM lactose sugar showing negligible staining for α2,6-Sial and α2,3-Sial levels. Cell are counter stained for nucleus with DAPI (white) and F-actin with Phalloidin (magenta). Scale bar: 100 μm.

**Figure S3:**
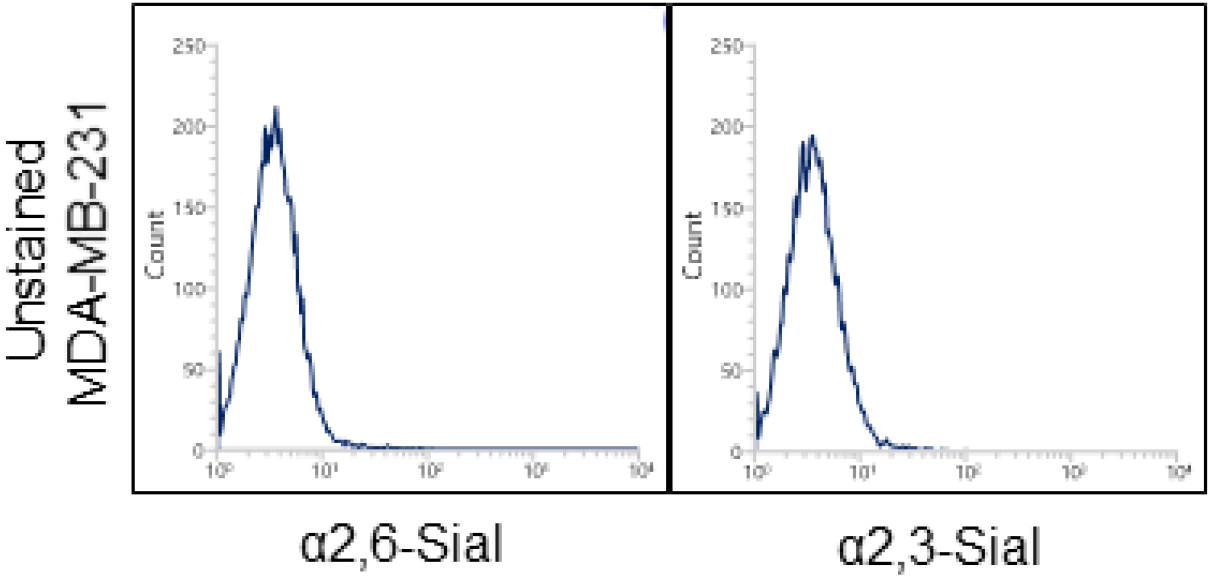
Lectin-based flow cytometry of unstained MDA-MB-231 cells showing basal auto fluorescence for α2,6- and α2,3-Sial levels.

**Figure S4:**
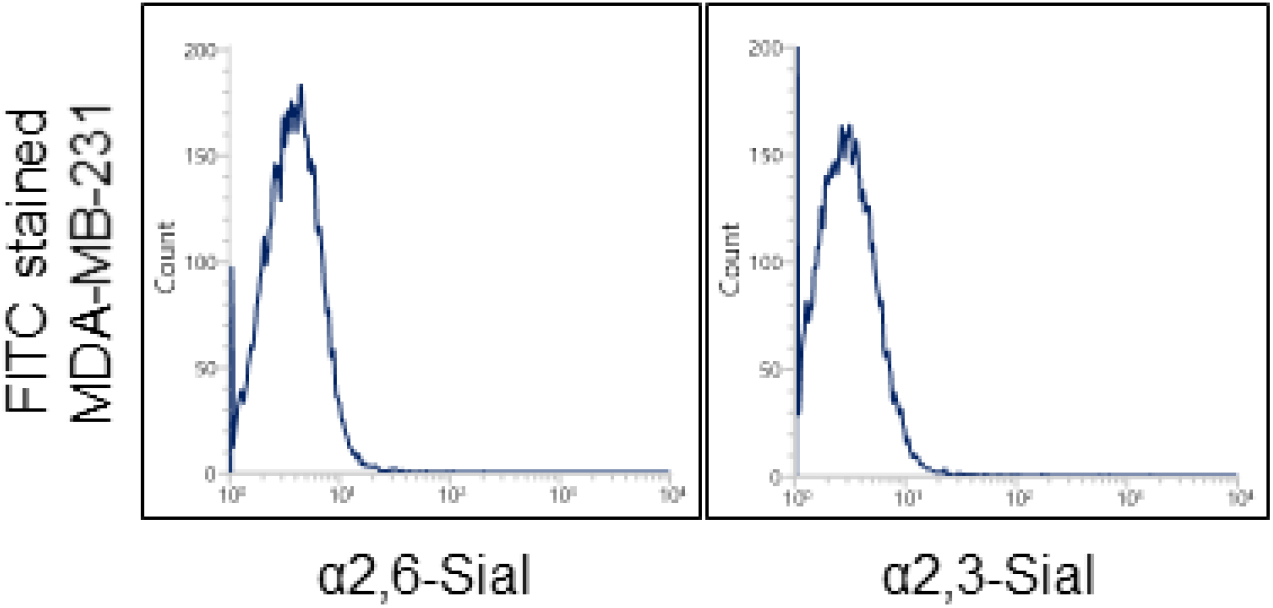
Lectin-based flow cytometry profiles of MDA-MB-231 cells showing basal auto fluorescence for α2,6- and α2,3-Sial levels upon staining with FITC fluorophore alone.

**Figure S5:**
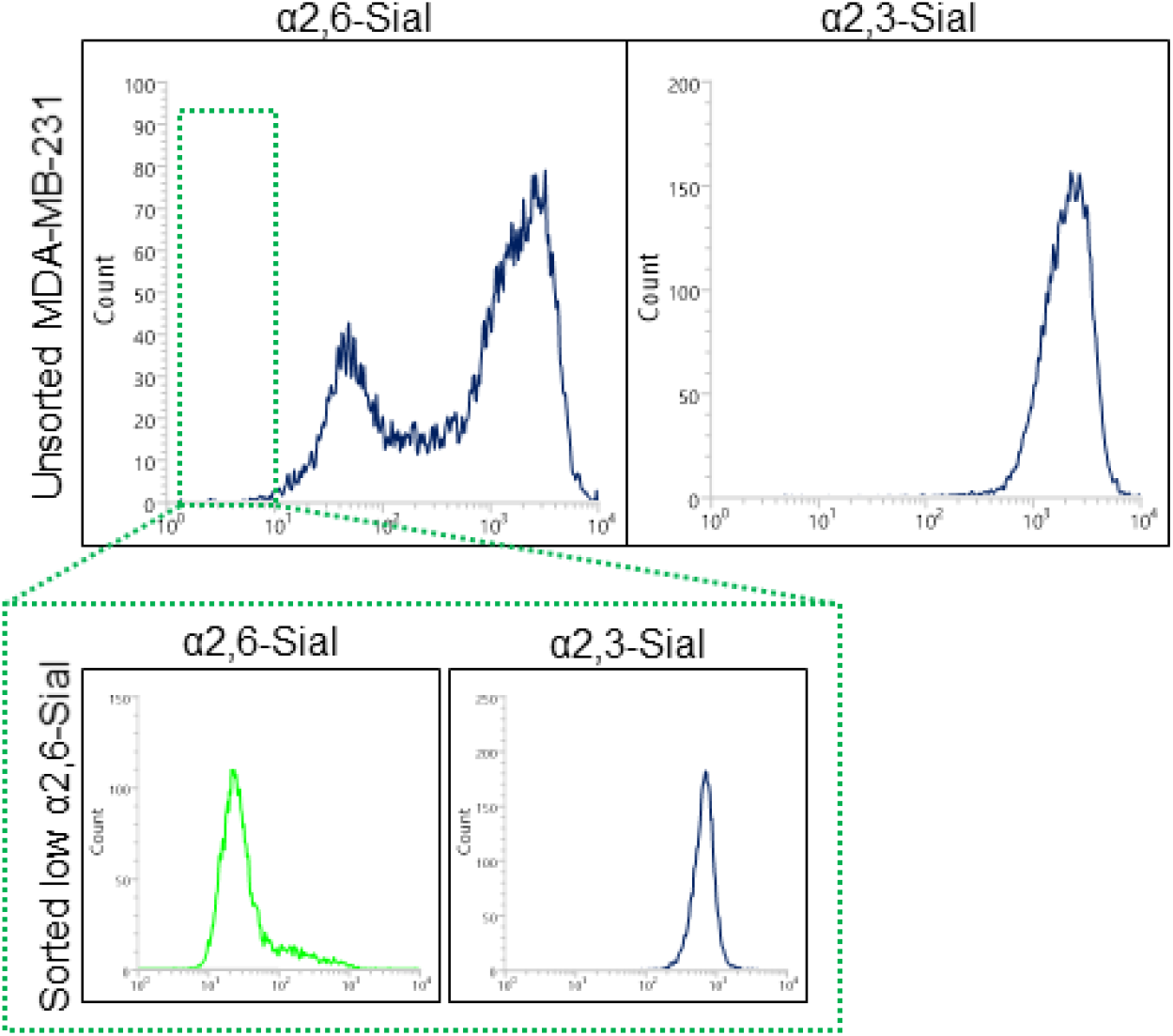
Lectin-based flow cytometry profiles of MDA-MB-231 cells showing bi-modal distribution of α-2,6 Sial levels on (top panel, left) and α-2,3 Sial levels showing uni-modal distribution (top right). Green inset shows moderate levels of α2,6-Sial (left) and unchanged α2,3-Sial levels in sorted low 2,6-Sial cells.

**Figure S6:**
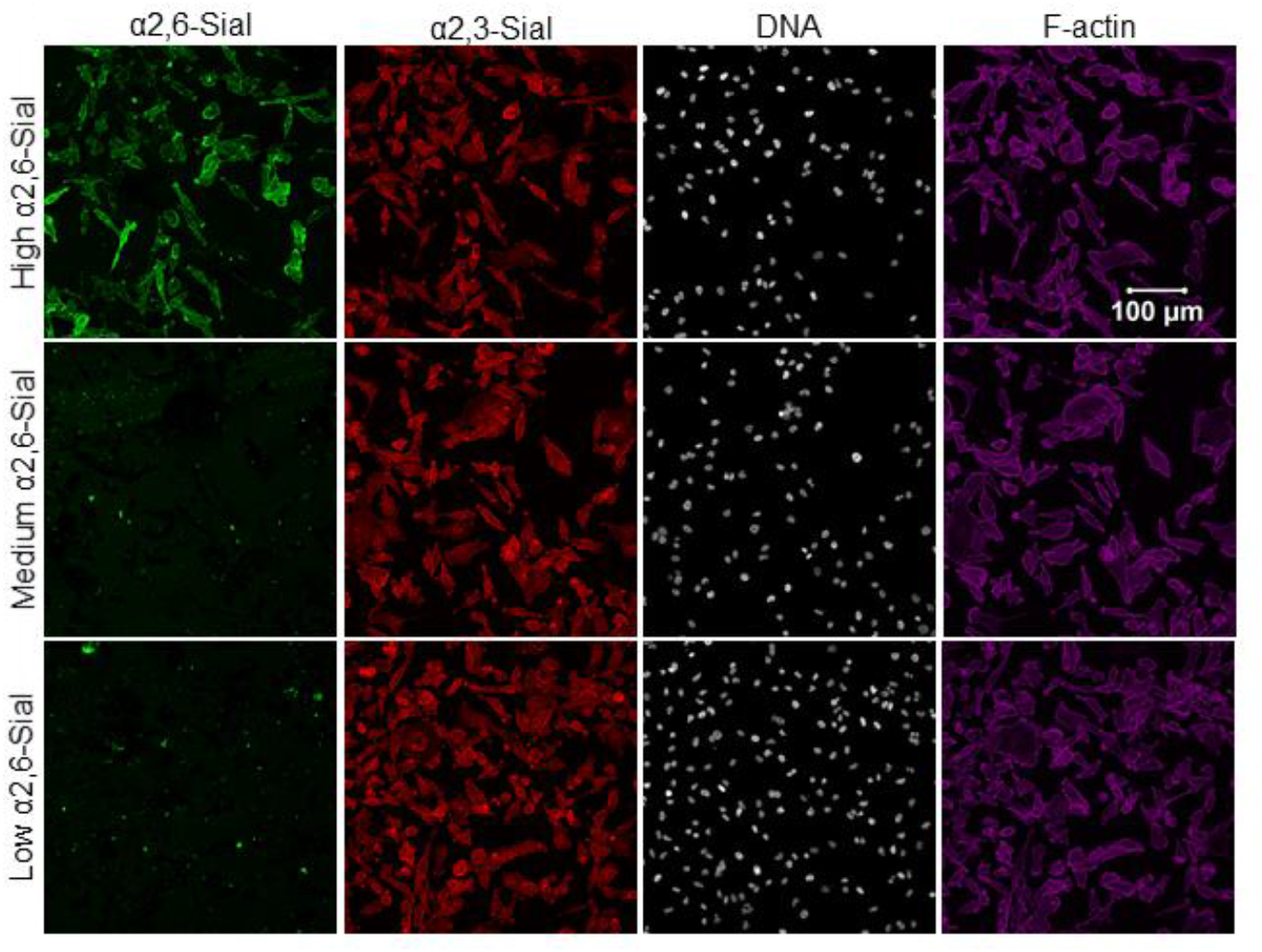
Confocal micrographs of sorted high-medium- and low 2,6-Sial cells stained for α2,6- and α2,3-Sial. High 2,6-Sial cells show highest staining for α2,6-Sial. Medium- and low 2,6-Sial cells show moderate staining for α2,6-Sial levels. All three populations show similar levels of α2,3-Sial levels. Cells are counter stained for nucleus with DAPI (white) and F-actin with Phalloidin (magenta). Scale bar: 100 μm.

**Figure S7:**
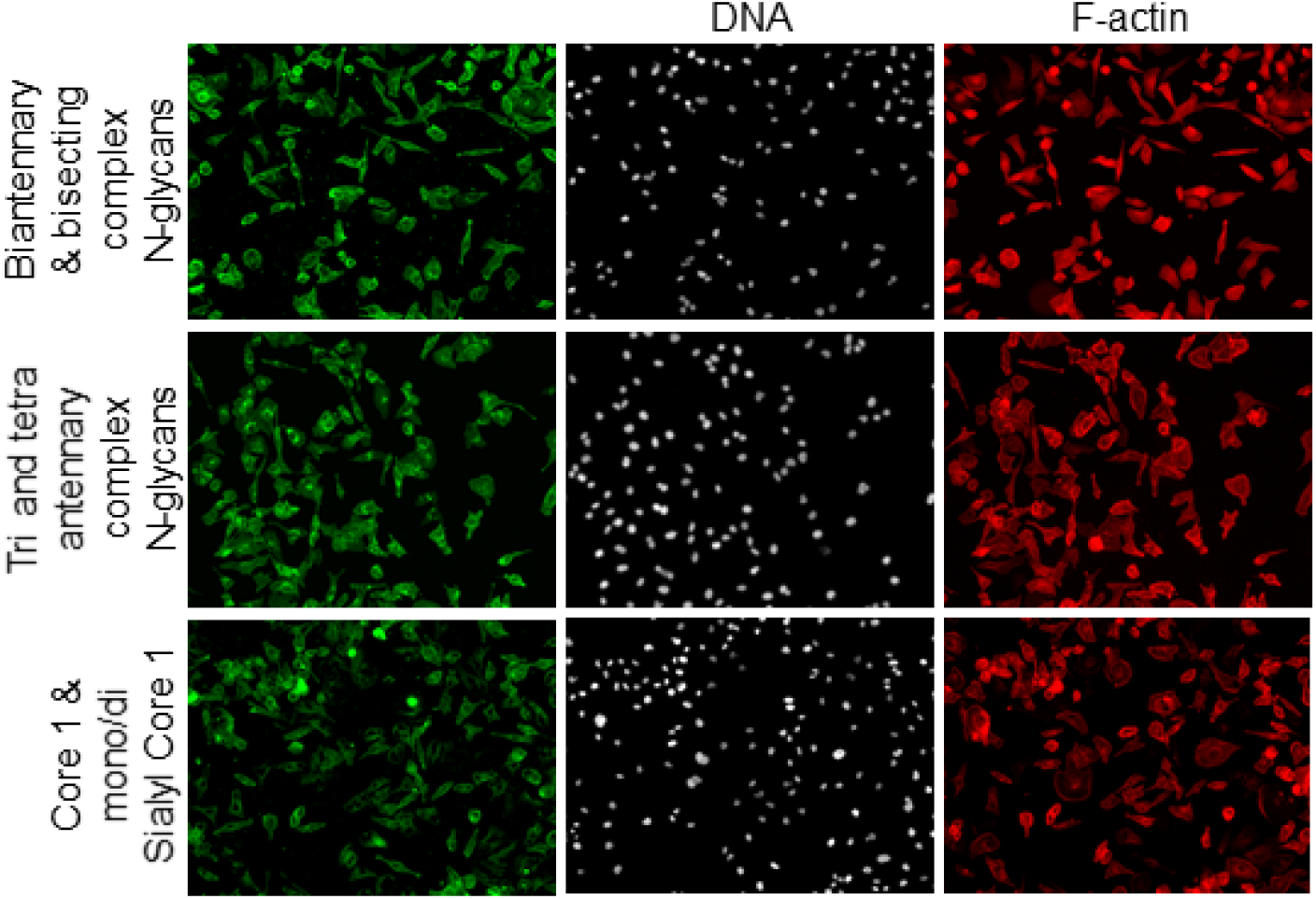
Epifluorescence micrographs of MDA-MB-231 cells showing uniform levels of bisecting & biantennary (top row, green), tri & tetra antennary (middle row, green) complex N-glycans and Core 1 & mono/di sialyl Core 1 (bottom row, green) O-glycans. Cells are counter stained for nucleus with DAPI (white) and F-actin with Phalloidin (magenta). Total magnification: 200X.

**Figure S8:**
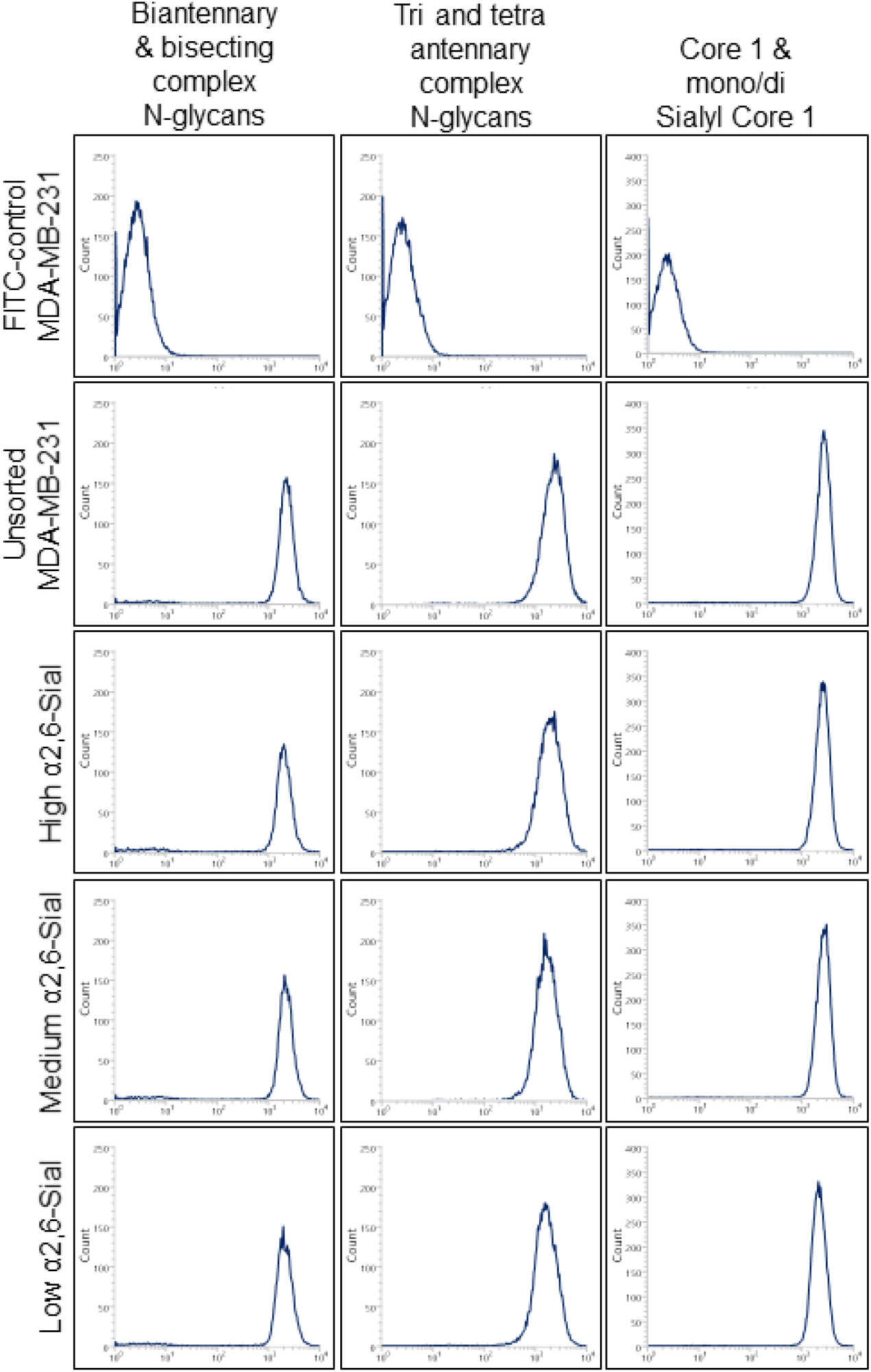
Lectin-based flow cytometry profiles of unsorted MDA-MB-231, sorted high-, medium- and low 2,6-Sial cells showing similar levels of bisecting & biantennary (left column), tri- and tetra antennary (middle column) complex N-glycans and Core 1 & mono/di- sialyl Core 1 O-glycans (right column).

**Figure S9:**
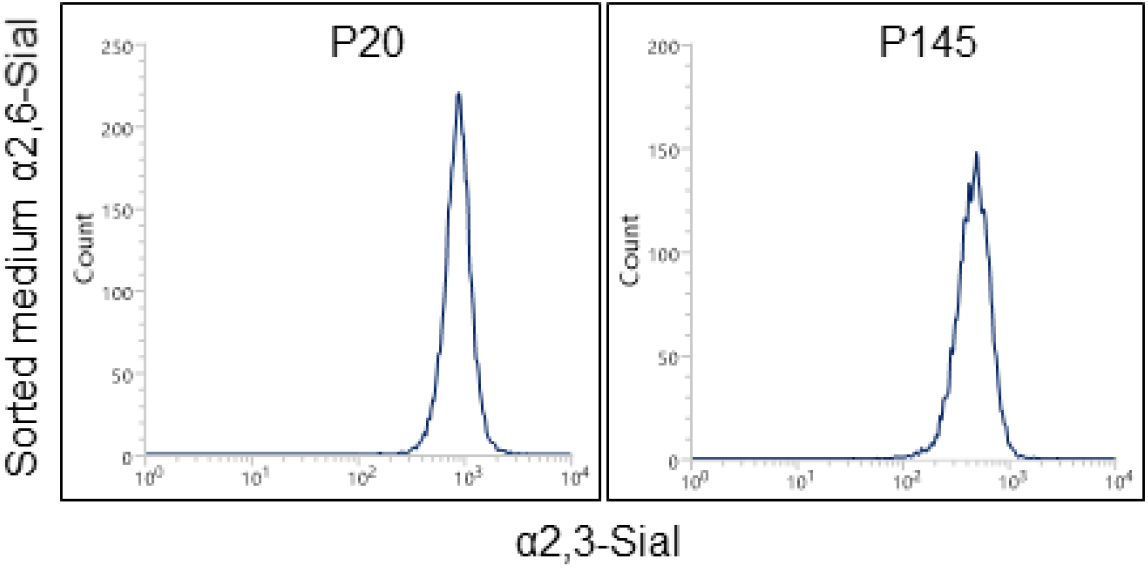
Lectin-based flow cytometry profiles of medium 2,6-Sial cells showing insignificant changes in α2,3- Sial levels at passage 20 and 145.

**Figure S10:**
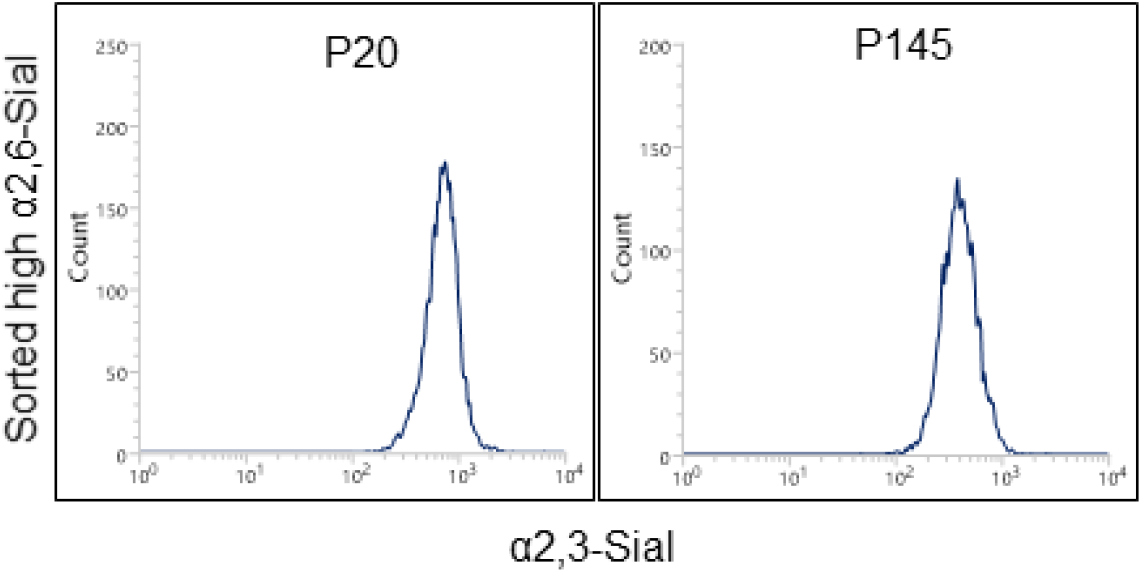
Lectin-based flow cytometry profiles of high 2,6-Sial cells showing insignificant changes in α2,3- Sial levels at passage 20 and 145.

**Figure S11:**
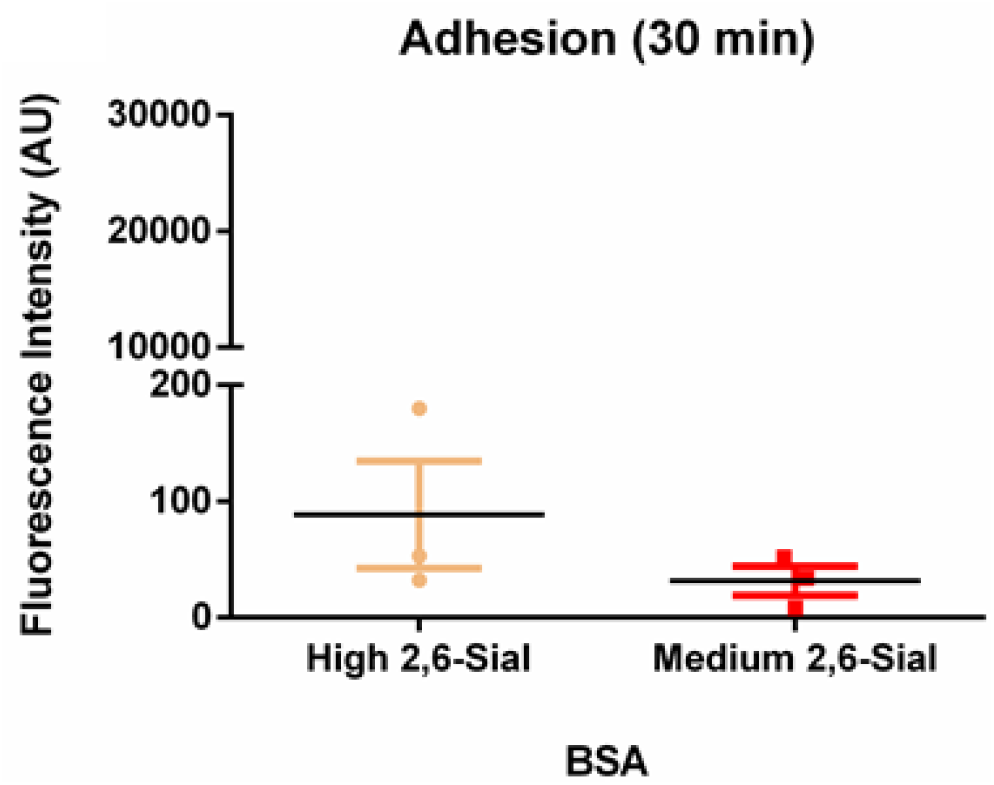
Graph showing negligible adhesion of high 2,6-Sial (yellow) and medium 2,6-Sial (red) cells to BSA coated plates.

**Figure S12:**
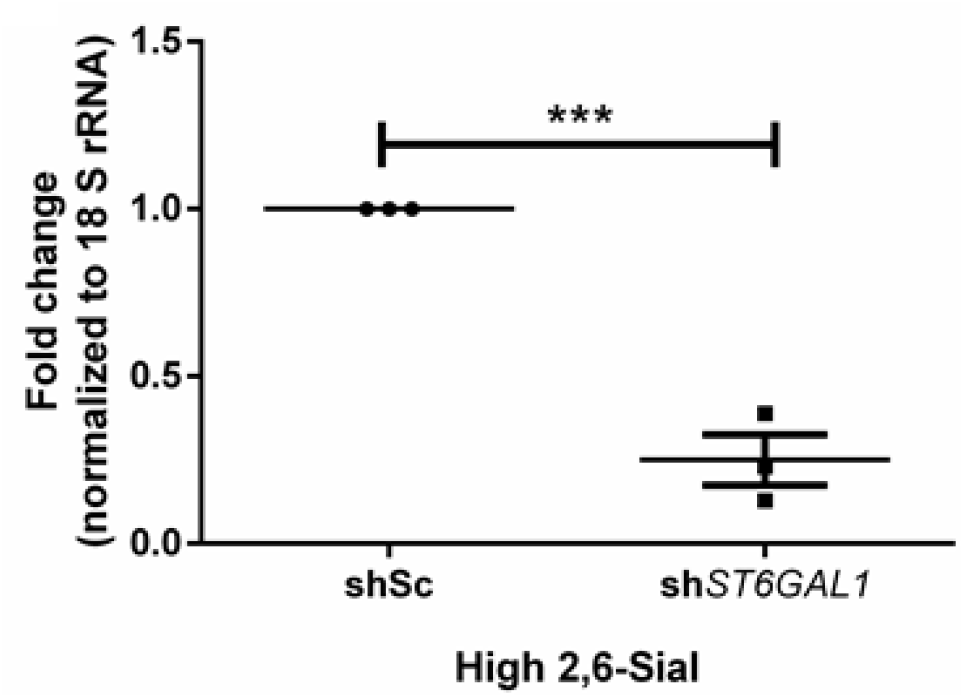
Graph showing significantly lower mRNA expression of *ST6GAL1* gene in sh*ST6GAL1* high 2,6- Sial cells when compared to shSc high 2,6-Sial control cells.

**Table S1:** Lectin list for glycan staining

| Sugar affinity | Lectin | Fluorephore conjugation |
| --- | --- | --- |
| α2,6-Sialic acid | *Sambucus nigra* | FITC |
| α2,3-Sialic acid | *Maackia amurensis* | TRITC |
| Bisecting/ bi-antennary complex N-glycan | *Phaseolus vulgaris lectin E* | FITC |
| Tri- & tetra-antennary complex N-glycan | *Phaseolus vulgaris lectin L* | FITC |
| Core-1 & mono/di Core-1 O-glycan | *Jacalin* | FITC |

**Table S2:**
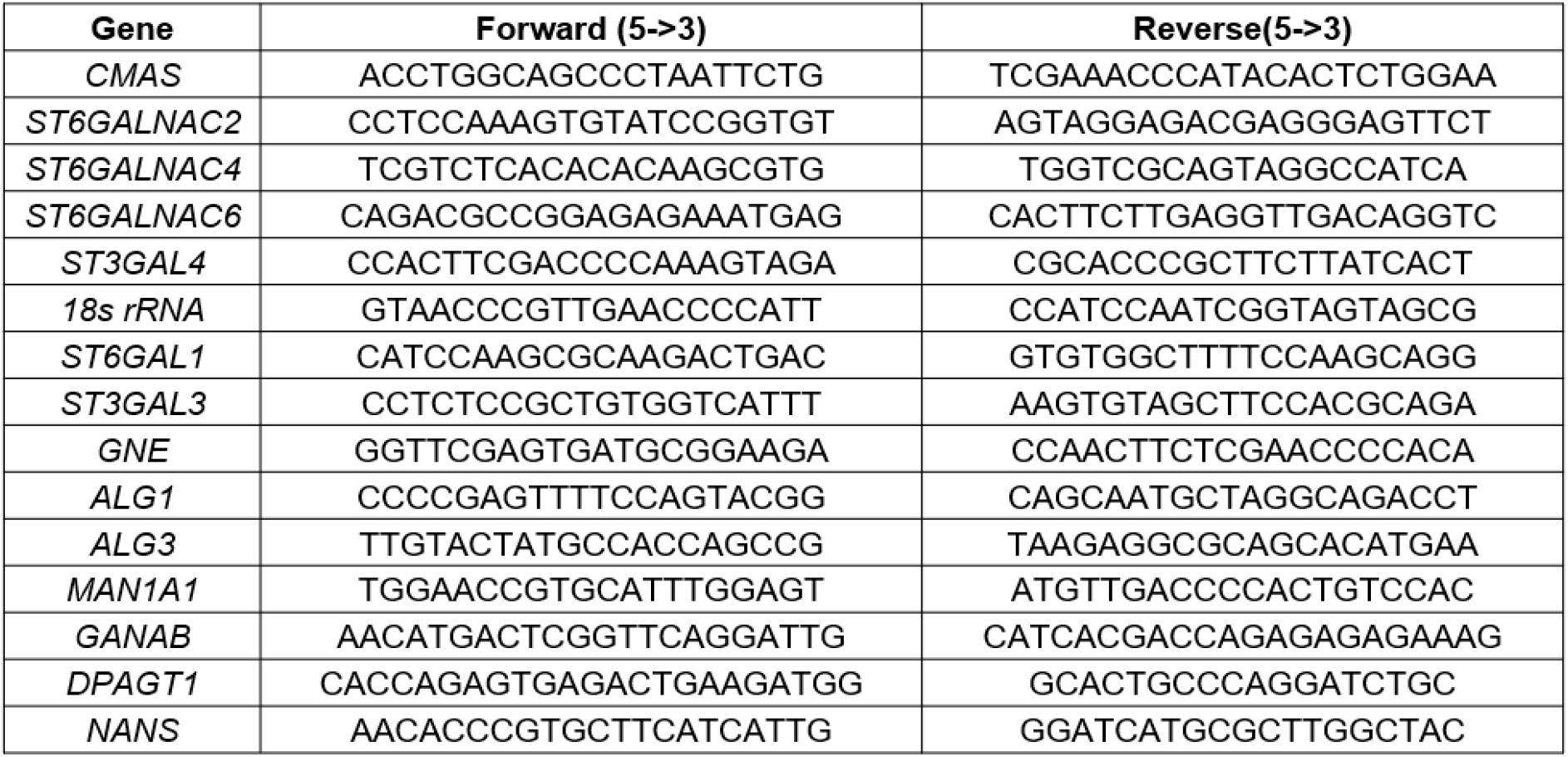
Primer list for qPCR

Video S1: Brightfield video of high 2,6-Sial cells invading into 3D fibrillar matrix. Images were taken from this video for figure 3C, top panel. Total magnification: 200X

Video S2: Brightfield video of medium 2,6-Sial cells invading into 3D fibrillar matrix. Images were taken from this video for figure 3C, bottom panel. Total magnification: 200X

Video S3: Brightfield video of shSc high 2,6-Sial cells invading into 3D fibrillar matrix. Images were taken from this video for figure 5B, top panel. Total magnification: 200X

Video S4: Brightfield video of *shST6GAL1* high 2,6-Sial cells invading into 3D fibrillar matrix. Images were taken from this video for figure 5B, bottom panel. Total magnification: 200X

